# Quantifying the Rearrangement Complexity of Pangenomes

**DOI:** 10.64898/2026.08.27.747493

**Authors:** Leonard Bohnenkämper, Jens Stoye

**Affiliations:** Bielefeld University, Faculty of Technology and Center for Biotechnology (CeBiTec)

## Abstract

The study of evolution between species (phylogenetics) and the study of evolution within a species (population genetics) are highly related, as the same biological mechanisms are fundamental to both fields. Although both have been studied for a long time, their joint study in a unified setting has been prevented by the different time scales they consider and the different data types they employ. A similar discrepancy holds for their whole-genome specializations, comparative genomics and pangenomics. Two active areas in these fields are genome rearrangement studies and graphical pangenomics, respectively. Since the emergence of graphical pangenomics, these have existed as separate fields, despite observations that central data structures representing genomic variants in both fields are highly similar.

While there exists a wealth of theoretical results for various rearrangement models in comparative genomics, the application to pangenomic data is hampered by the limitations of rearrangement problem formulations. On the practical side, pangenomes typically contain too many individual genomes for classical problems, such as the often NP-hard parsimony problems, to be solved, or for all-vs-all comparisons using rearrangement distances to be performed. On the theoretical side, some assumptions in the formulation of rearrangement problems, such as the assumption of an underlying tree, are inadequate for many pangenomes.

In this work, we propose the *Complete Ancestral Reconstruction for Pangenomes (CARP)* problem, which overcomes these limitations while retaining intuitive relationships to both classical rearrangement problems and pangenome graphs.

## 1 Introduction

Rearrangements of genetic material have been described even before the molecular structure of DNA was known [44, 4, 15]. These rearrangements and other structural variations have been discovered not only to be ubiquitous, but also to play a significant role in evolution [49, 16, 19], as well as in genetic diseases like cancer [7, 43, 3, 29]. Four years before the first whole eukaryotic genome was sequenced, David Sankoff proposed a framework to computationally quantify rearrangements between two genomes, the rearrangement distance problem [42]. Thirty years later, this basic distance problem has been solved for many different types of rearrangement models, some with simple results, like the Single-Cut-or-Join (SCJ), Double-Cut-and-Join (DCJ) and block interchange models [8, 18, 51], and some with more involved results, like the Reversal, Hannenhalli-Pevzner (HP) and DCJ-indel models [6, 21, 22], to name a few.

In the intervening decades, advances in sequencing technology have greatly enhanced the available material for analysis. This enabled the simultaneous study of several genomes in a clade – the field of pangenomics emerged. With advances in assembly quality, pangenomics itself moved from studying collections of genes [46] to collections of sequences [5] to studying entire genomes [47]. These sets of genomes are typically compressed for analysis, either via lossless text indices such as the Burrows Wheeler Transformation [41, 52], or via lossy graphs such as colored de Bruijn [34, 26, 10] or variation graphs [20, 17]. There exist many different ways to quantitatively characterize pangenomes, such as openness [46, 37] or core percentage [25]. Many of these are already possible using gene-based pangenomes. One truly novel possibility, now that pangenomes are constructed from complete chromosomes, is the quantification of their structural complexity.

Examining the graphical representations of pangenomes, quantifying this complexity is intuitively possible. For example, in Fig. 1, one can hardly deny that Graph B is structurally more complex than Graph A. Of course, an objective, numerical measure is more desirable, especially in less obvious instances.

**Figure 1:**
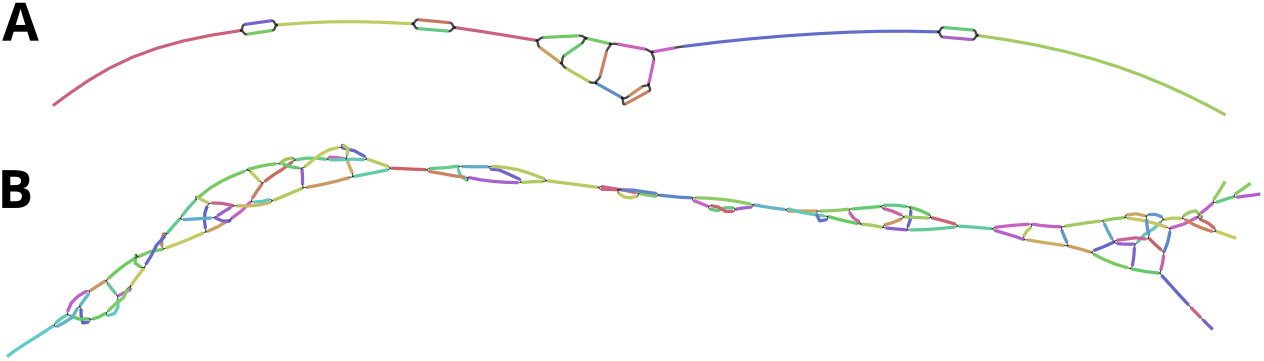
1000 BP regions in two compacted de Bruijn graphs (*k* = 31) built with Bifrost [26] and visualized with Bandage [48]. A - built from 56 *Yersinia pestis* genomes. B - built from 48 *Escherichia coli* genomes.

**Figure 2:**
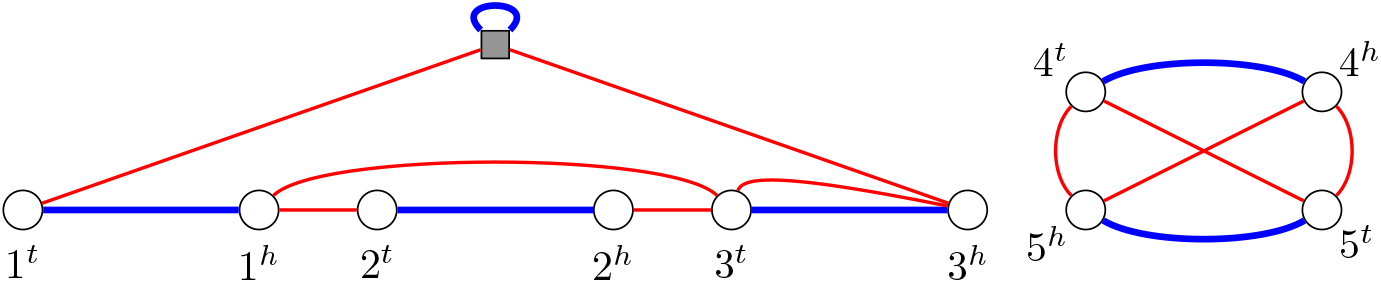
Multiple Breakpoint Graph (MBG) for the pangenome 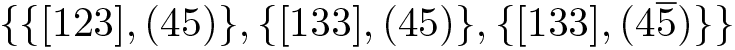. Blue edges are marker edges, red edges are adjacency edges, and the quadratic grey node is the telomere Ø.

If one were to consider a “pangenome” of exactly 2 genomes, classical rearrangement analyses already give an answer to this quantification, namely the aforementioned rearrangement distance under a certain model. Notably, the rearrangement distance across various models can be quantified using the *breakpoint graph* [21, 22, 51, 9]. This graph bears remarkable similarities to pangenome graphs. Specifically the close relationship between breakpoint and de-Bruijn graphs has been examined in [1] and [32].

Interestingly, the breakpoint graph of two identical genomes, whose distance is therefore 0 under classical rearrangement models, corresponds to a graph without any branching nodes, as seen in the bottom graph in Fig. 3. A pangenome with a graph like this has arguably minimal rearrangement complexity. Therefore, we say such a graph and any pangenome with such a graph is *simple*.

**Figure 3:**
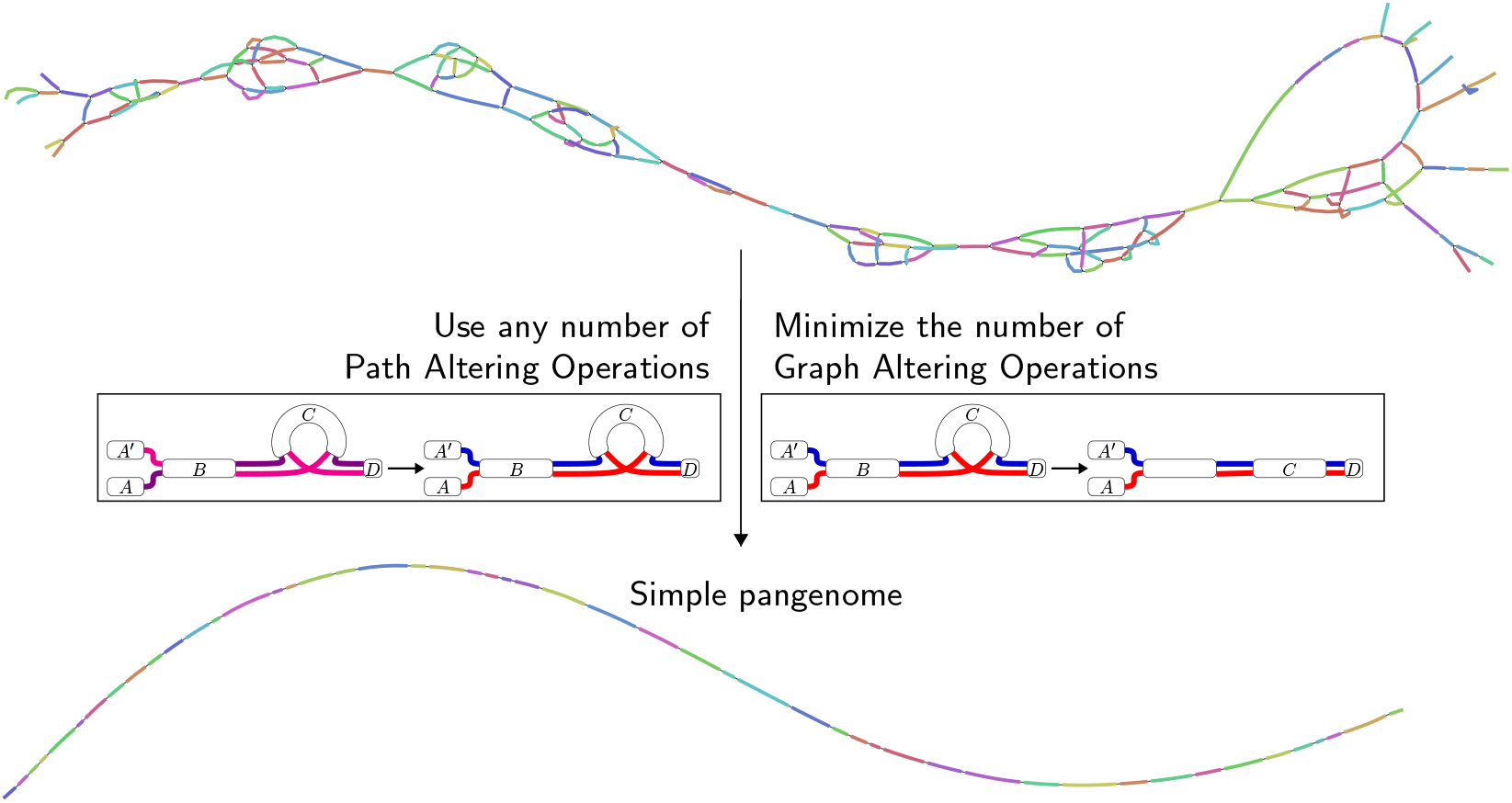
Schematic representation of the CARP rearrangement problem: A pangenome is transformed into the closest simple pangenome using the minimum number of Graph Altering Operations while utilizing arbitrarily many operations from a complete set of Path Altering Operations. An example of a Path Altering Operation (a homologous recombination between the chromosome 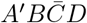 (pink) and the chromosome *ABCD* (purple) at marker *B*, transforming them into *A*′*BCD* (blue) and 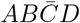 (red)) is shown in the left box. An example of a Graph Altering Operation (a reversal of marker *C* in the chromosome 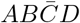 (red)) is shown in the right box. Note that both types of operations change paths directly, but only Graph Altering Operations change edges in the graph.

In this work, we propose the *Complete Ancestral Reconstruction for Pangenomes (CARP)* problem (Section 2), a new rearrangement problem designed to quantify the structural complexity of a pangenome based on how many graph altering operations, under a given model, it takes to transform the pangenome into a simple pangenome (see also Fig. 3). We then show how this problem relates to classical rearrangement problems (Section 3). We give a solution to this problem for the SCJ model (Section 5) and demonstrate in a number of experiments (Section 6) how even under this simple model, CARP can be used to derive valuable insights about rearrangements in pangenomes and pangenome graphs. This includes comparative analyses of the rearrangement complexities between different pangenomes and different pangenome graphs constructed from the same sequence set. We further discuss the potential of the CARP problem formulation (Section 7), particularly once solutions for more advanced rearrangement models become available.

## 2 A new rearrangement problem

In a pangenome graph, each node represents a sequence segment. We call these segments *markers*. An edge connecting two nodes represents an *adjacency*, meaning that these two markers are immediate neighbors on a chromosome. To that end, an edge can either connect to the beginning or to the end of a marker. Each chromosome is then represented as a path or a cycle in this graph, depending on whether it is linear or circular.

A notable difference between pangenome and breakpoint graphs is that the latter aim to represent only rearrangements, and not local variations such as SNVs or small indels. To capture only structural variations, one can use genes or *collinear blocks* as markers. Collinear blocks can be generated as a by-product of whole genome alignment, such as in Mummer [14] or Mauve [11, 12], but some pangenome graph construction methods, such as Minigraph-Cactus [23] or Pangraph [35], are also able to output collinear blocks. Additionally, there is dedicated software for collinear block detection, such as SibeliaZ [33].

What one considers a rearrangement, is to some degree dependent on a size threshold – note that one could even consider the deletion of a single base a “structural” variation. We will therefore presume that, using one of the aforementioned methods, a suitable level of abstraction that defines markers has been found.

Formally, a marker is a symbol from a large alphabet Σ. A *chromosome* is then associated with a string of oriented markers 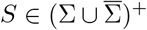 where 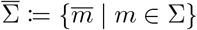 and 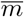 denotes marker *m* in reverse orientation. We call the string *S* a *chromosome string*. A convenient way of writing a marker *m* in this context is by referring to its *extremities*, that is, to its beginning *m*^*t*^ (*tail*) and to its end *m*^*h*^ (*head*). Then *m* = *m*^*t*^*m*^*h*^ and 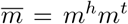. Two neighboring markers *m* and *n* in a string induce a pair of neighboring extremities, an *adjacency*. Depending on their relative orientation, adjacencies are *m*^*h*^*n*^*t*^, *m*^*t*^*n*^*t*^, *m*^*t*^*n*^*h*^ and *m*^*h*^*n*^*h*^ for *mn*, 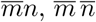 and 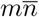, respectively. The identity of an adjacency is purely determined by the involved extremities, not their order, i.e. *m*^*h*^*n*^*t*^ is the same adjacency as *n*^*t*^*m*^*h*^.

A chromosome may be *linear*, in which case we denote it by [*S*], or circular, in which case we denote it by (*S*). A circular chromosome (*s*_1_ … *s*_l_) in addition to the adjacencies of *s*_1_ … *s*_l_ has the additional adjacency induced by *s*_l_*s*_1_. For a linear chromosome [*S*] it is helpful to define the additional adjacencies ∅*u* and *v*∅ for the extremities *u* and *v* with which the chromosome starts and ends. We call ∅ a *telomere*. Note that we regard chromosomes as equal even when denoted in reverse orientation, i.e. 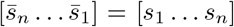 and circular chromosomes as equal even when denoted in different orientation (*s*_k_ … *s*_n_*s*_1_ … *s*_k−1_) = (*s*_1_ … *s*_k−1_*s*_k_ … *s*_n_). A *genome* is a multiset of chromosomes, and a *pangenome* is a multiset of genomes.

We can now define the pangenome graph that represents the order of these markers as a bi-edged graph [38]. For ease of notation, we write the (undirected) edge between vertices *u* and *v* as *uv* or *vu*.

### Definition 1.

*The* Multiple Breakpoint Graph (MBG) *G*(ℙ) *of a pangenome* ℙ *is an undirected graph* (*V, E*_*adj*_ ∪ *E*_*mark*_) *where V represents the extremities of* ℙ *including the telomere node* ∅ *and E*_*adj*_, *E*_*mark*_ *are two types of edges:*

- *E*_*adj*_ *is a set of undirected edges, which is the union of all adjacencies in* P. *We call E*_*adj*_ adjacency edges.
- *E*_*mark*_ *is a set of undirected edges that contains the edge m*^*t*^*m*^*h*^ *for each marker m in addition to a self edge* ∅∅ *of the telomere* ∅. *We call E*_*mark*_ marker edges.

In principle, any graph in `gfa` format can be represented as a MBG if telomeres are added to those nodes at the end of linear paths. However, note that the theory presented here assumes all variants represented in the graph are captured by the rearrangement model used. An example of a MBG is found in Fig. 2.

Recall the earlier motivation of a *simple* pangenome without any branching paths in its pangenome graph as having minimal structural complexity. We can now formulate this more precisely. A pangenome and its MBG are *simple* if for each marker *m* (excluding ∅) at most one adjacency edge connects to its beginning *m*^*t*^ and at most one adjacency edge connects to its end *m*^*h*^. We then wish to quantify the structural complexity of a pangenome by the number of rearrangement steps necessary to transform it into a simple pangenome. The operations used for this transformation are most meaningful when modeled after biological events, such as reversals or recombinations that occur in chromosomes of the pangenome. They must thus fundamentally operate on chromosomes, i.e. the paths in the graph, not the graph itself. However, not all operations have the same effect on the MBG. There are operations that never affect the structure of the MBG, such as the duplication of a chromosome or the recombination of two chromosomes, as shown in the left box of Fig. 3. Other operations, typically ones that are associated with breaking a chromosome, may have an effect on the graph. Such an operation is shown in the right box of Fig. 3. We call these types of operations Path Altering Operations and Graph Altering Operations, respectively. Formally, an operation *o* is a *Path Altering Operation (PAO)* if, after applying it to a pangenome ℙ, denoted 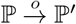, the resulting pangenome ℙ′ has the same MBG as ℙ, i.e. *G*(ℙ′) = (ℙ). Otherwise it is a *Graph Altering Operation (GAO)*.

Classical rearrangement operations, such as reversal, translocation, block interchange and DCJ, are all GAOs. In fact, one can reformulate many classical problems as transforming a set of input genomes into the closest set of genomes with a simple MBG by using the model operation and a limited set of supplementary PAOs while minimizing the number of model operations used. We examine this relationship for the distance, median, large parsimony and halving problems in Section 3.2.

However, in these formulations the interaction between lineages is strictly limited – owing to the fact that most classical rearrangement problems (implicitly) assume an underlying tree. In fact, the only PAOs necessary to model these classical problems are coalescence (unifying equal genomes) and some form of chromosome de-duplication (unifying equal chromosomes in the same genome). In a pangenome, however, the interactions between lineages can be much more frequent and complex, such as the transfer of plasmids between bacteria or the sexual reproduction of many eukaryotes. We therefore propose to use a “maximally powerful” set of PAOs to quantify the rearrangement complexity of a pangenome. We call such a set *complete*. Formally, we say a set of PAOs is *complete* if it is able to transform any two pangenomes with the same MBG into each other. Such a set exists; we give it in Theorem 1 (Section 4.3).

The definition for the *Complete Ancestral Reconstruction for Pangenomes (CARP)* problem is then as follows: Given a pangenome, a set of GAOs and a complete set of PAOs, find a simple pangenome into which it can be transformed using any number of PAOs from a complete set and a minimal number of GAOs. A visual summary is found in Fig. 3. Formally, we define the problem as follows.

**Problem 1** (Complete Ancestral Reconstruction for Pangenomes (CARP)). *Given a pangenome* ℙ, *a complete set of PAOs N and a set of GAOs A, find the shortest sequence of GAOs o*_1_ … *o*_n_ ∈ *A*^*^, *such that there is a sequence of operations s* = *O*_0_*o*_1_*O*_1_ … *o*_n_*O*_n_ *transforming* ℙ *into a simple pangenome* A *where each O*_i_ *is a (possibly empty) sequence of PAOs, O*_i_ ∈ *N* ^*^. *We call the simple pangenome* A *a* CARP simplification, *s a* CARP scenario *and n the* CARP measure *of* ℙ.

## 3 Relationship to classical rearrangement problems

CARP is related to classical rearrangement problems, such as the distance, median and large parsimony problem. In fact, all of these can be expressed as specializations of a common superproblem, the *General Ancestral Reconstruction Problem*, which we examine in Section 3.2. In essence, these problems differ only by which path altering operations they permit. We thus first examine a few path altering operations in Section 3.1. Finally, we discuss a complete set of path altering operations in Section 4.3.

### 3.1 Path Altering Operations relevant to classical problem formulations

In this section, we give examples of PAOs that are useful to understand the relationship between CARP and classical rearrangement problems (see Section 3.2). One of the most fundamental PAOs is the emergence of new lineages, which when regarded backwards in time is known as *coalescence*. In the following, we use multi-set addition ⊕ and multi-set subtraction ⊖.

#### Definition 2

(Coalescence/Divergence). *Given a pangenome* ℙ *and genomes G*_1_, *G*_2_, …, *G*_k_ ∈ ℙ *with G*_1_ = *G*_2_ = … = *G*_k_, *a k*-coalescence *on genomes G*_1_, *G*_2_, …, *G*_k_ *creates the pangenome* ℙ′ = ℙ ⊖ {*G*_2_, …, *G*_k_}. *Given a pangenome* ℙ *and a genome G*_1_ ∈ P, *a k*-divergence *on genome G*_1_ *creates the pangenome* ℙ′ = ℙ ⊕ {*G*_2_, …, *G*_k_} *with G*_2_ = … = *G*_k_ = *G*_1_.

As we examine in Section 3.2, many classical problems examine only tree-like evolution without duplicated markers and can thus be formulated directly using only coalescence. However, some classical problems, such as double distance or halving problems, require the modeling of whole genome duplications. To that end, we first define the duplication and de-duplication of a single chromosome.

#### Definition 3.

*A k*-multiplication (→*) transforms a single chromosome in one of the following ways and a k*-de-multiplication (←*) transforms either k identical or one repetitive circular chromosome in one of the following ways:*

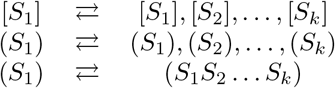

*where S*_2_ = … = *S*_k_ = *S*_1_.

A whole genome duplication is then characterized by duplicating all of a genome’s chromosomes. Again, there is an inverse when viewing time “backwards”.

#### Definition 4.

*A* genome duplication operation *transforms a genome by* 2*-multiplying all of its chromosomes. A* genome halving operation *is the inverse of a genome duplication operation*.

These definitions suffice to characterize many classical rearrangement problems as we examine in the following section.

### 3.2 The General Ancestral Reconstruction Problem (GARP)

Under the assumption that PAOs do not create new structural complexity while a simple set of genomes possess minimal structural complexity, we can quantify the complexity of a set of genomes by measuring how many GAOs it takes to transform it into a simple pangenome while using some supplementary set of PAOs. This formulation (see Problem 2) is then a common framework for CARP and classical rearrangement problems.

**Problem 2** (General Ancestral Reconstruction Problem (GARP)). *Given a pangenome* ℙ, *a set of PAOs N and a set of GAOs A, find the shortest sequence of GAOs o*_1_ … *o*_n_ ∈ *A*^*^, *such that there is a sequence of operations s* = *O*_0_*o*_1_*O*_1_ …

… *o*_n_*O*_n_ *transforming* ℙ *into a simple pangenome* A *where each O*_i_ *is a (possibly empty) sequence of PAOs, O*_i_ ∈ *N* ^*^.

Note that CARP (Problem 1) is a specialization of Problem 2, where the set of PAOs is required to be *complete*. In classical rearrangement problems, this is not the case.

For a simple example from classical rearrangement literature, consider the genomic distance problem [42]. In this problem, two genomes *X* and *Y* are given that contain the same set of markers, each exactly once. The goal is to find the minimum number of rearrangements to transform *X* into *Y* under a given model *M* . If each rearrangement operation in *M* has an inverse, which is typically the case, we can model this in the GARP framework.

We leave the set of PAOs *N* empty and define the set of GAOs *A* as the operations of *M* . Then for each scenario transforming *X* into *Y*, 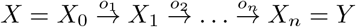, there is a GARP solution with the same number of GAOs, namely *o*_1_*o*_2_ … *o*_n_. Note that two genomes without duplicate markers are equal if and only if their adjacencies are equal [2], meaning the MBG constructed from both genomes is simple. Since *X*_n_ = *Y*, the final MBG is simple. Thus, the number of GAOs required in the GARP formulation is a lower bound for the distance.

Conversely, the distance also bounds the number of GAOs in the GARP scenario. To see why, we can find a classical rearrangement scenario for any GARP scenario. In this case, the scenario can modify both *X* and *Y* with 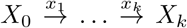 and 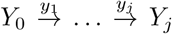. Because *G*({*X*_*k*_, *Y*_*j*_ }) is simple, *X*_k_ = *Y*_j_. Thus, we can use the inverse operations 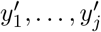 of *y*_1_, …, *y*_j_ respectively to build the classical scenario 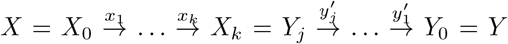 . The problem formulations are thus equivalent.

A slightly more involved example is the large parsimony problem [2] (called “Multiple Genome Rearrangement Problem” there). In this problem, extant genomes *X*_1_, …, *X*_n_ are given and one is asked to find a binary tree *T* with *X*_1_, …, *X*_n_ at the leaves and genomes *Y*_1_, …, *Y*_n−1_ at the internal nodes, such that the total distance along the edges *E*(*T*), ∑_(U,V)∈E(T)_ *d*(*U, V*), is minimized. Under the assumption that the operations of *M* are symmetric, the following GARP formulation is equivalent: In Problem 2, let ℙ = *X*_1_, …, *X*_n_, let *N* contain only 2-coalescence operations, and let *A* be the operations of *M* . We show how to obtain a GARP solution from the large parsimony solution. Assume the children *V, W* of an internal node *U* exist in the current pangenome. From the large parsimony solution we can deduce operations *v*_1_ … *v*_k_ transforming *U* into *V* and operations *w*_1_ … *w*_j_ transforming *U* into *W* . Since operations have an inverse, there are operations 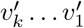 and 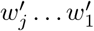 transforming *V* and *W* into *U*, which we can use in the GARP scenario and finally use a 2-coalescence to unify the genomes into *U* . We can apply this procedure bottom up to arrive at the corresponding GARP scenario. Conversely, one can find the corresponding large parsimony problem solution for a GARP scenario by using the genomes unified by the 2-coalescence as inner nodes and reversing the scenarios to be top-down. In fact, this is how Alekseyev and Pevzner constructed the MGRA heuristic for the large parsimony problem for 2-breaks (DCJ) in [2], although they do not explicitly use coalescence, but instead rely on the concept of “strict transformations”, i.e. such transformations that conform to a phylogeny.

In the same way one can find a GARP formulation for many classical rearrangement problems. We give a selection in Table 1. We note that coalescence is the only interaction between genomes in these problems. As mentioned earlier, due to the phylogenetic closeness of the genomes in a pangenome, the interactions in a pangenomic context may well be more frequent and direct. For our pangenome focused specialization of GARP, CARP, we permit much more interaction by using a complete set of PAOs. We give one example of such a complete set in the following section.

**Table 1:**
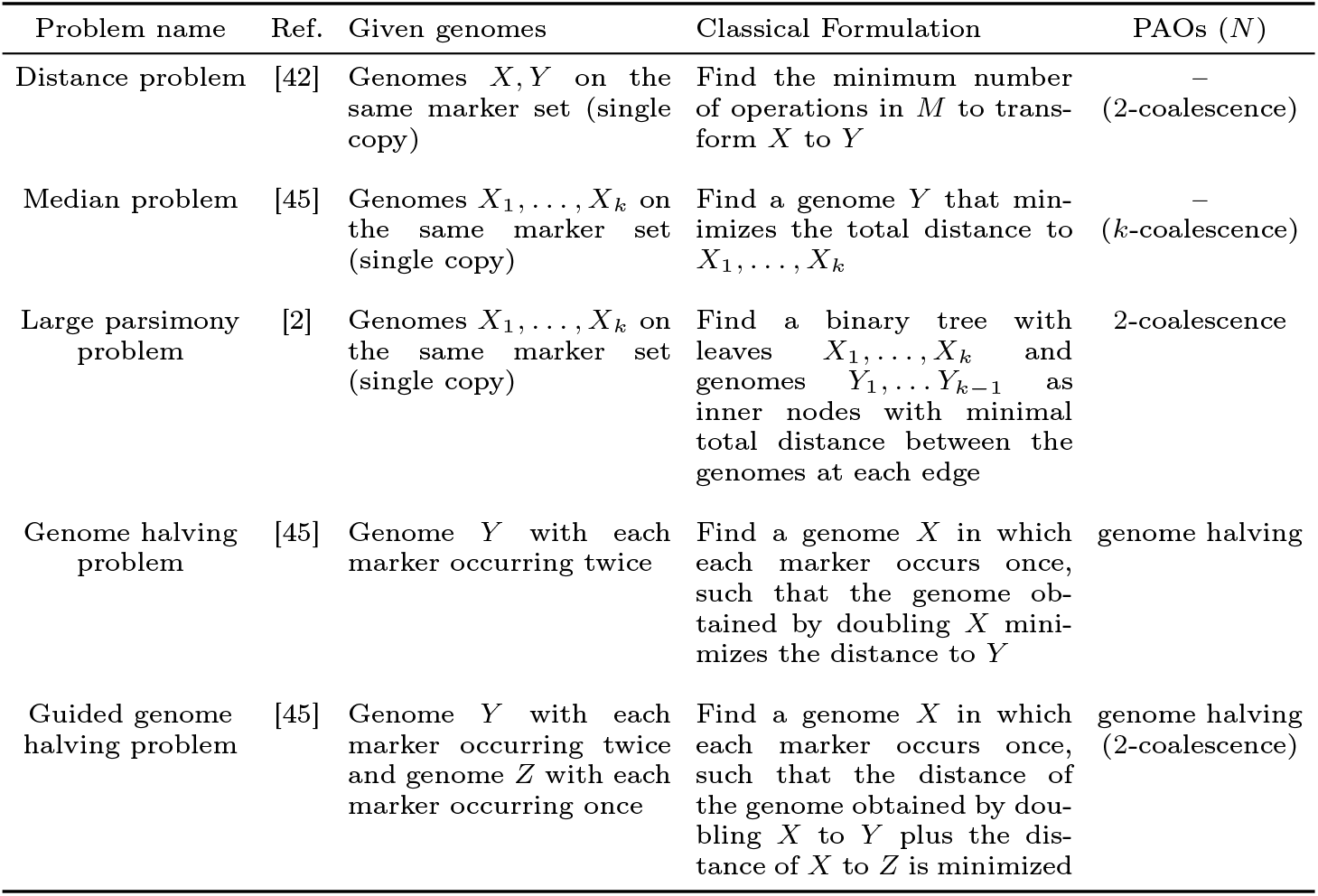
Classical genome rearrangement problems on a model *M* with symmetric operations and an equivalent GARP problem (see Problem 2). A dash (–) represents that no PAOs are needed while brackets indicate those PAOs that one can include into the problem formulation without losing the equivalence. In each case the set of GAOs *A* equals the set of operations of *M* and P is the multiset containing all input genomes.

**Table 2:**
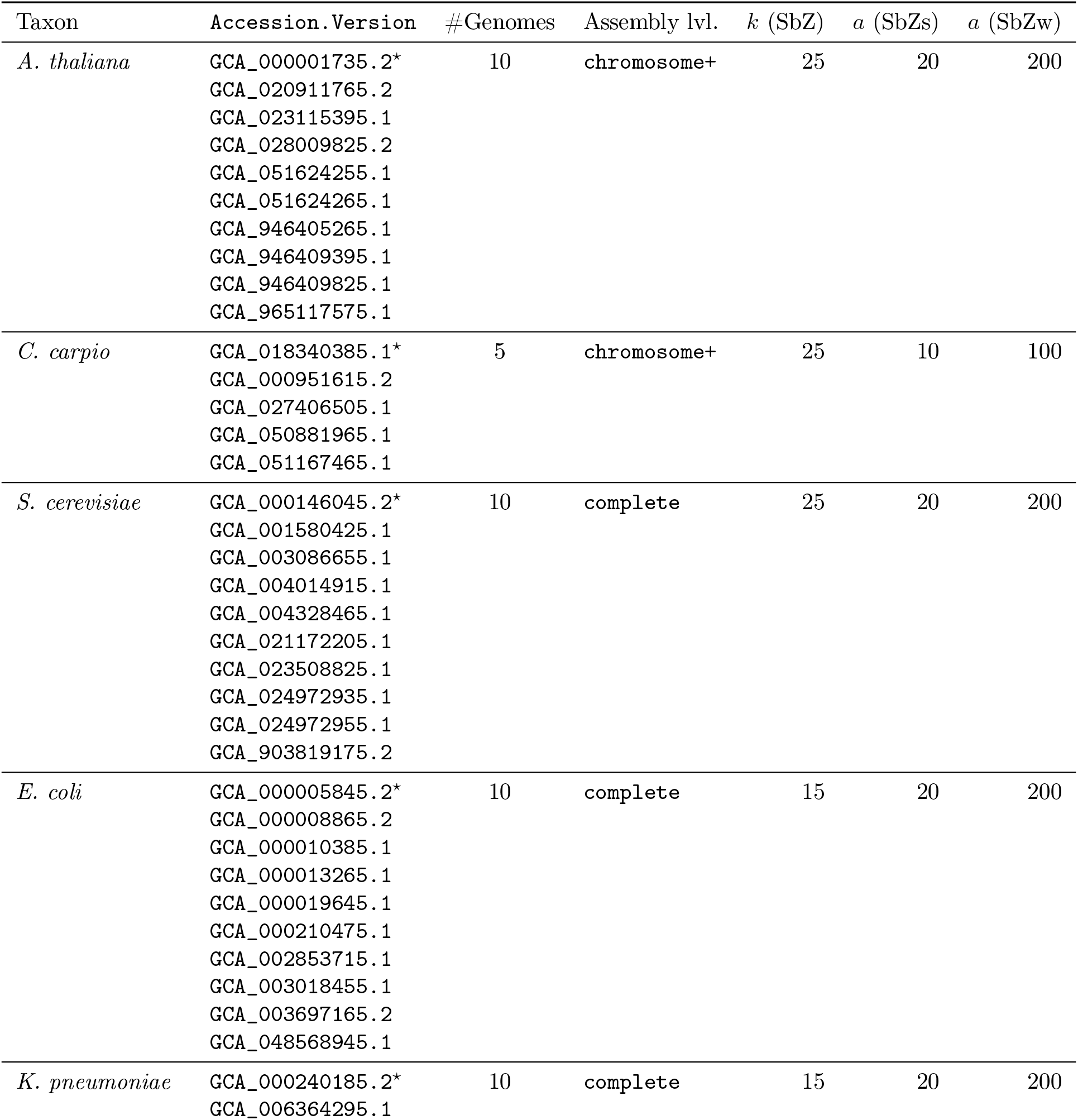
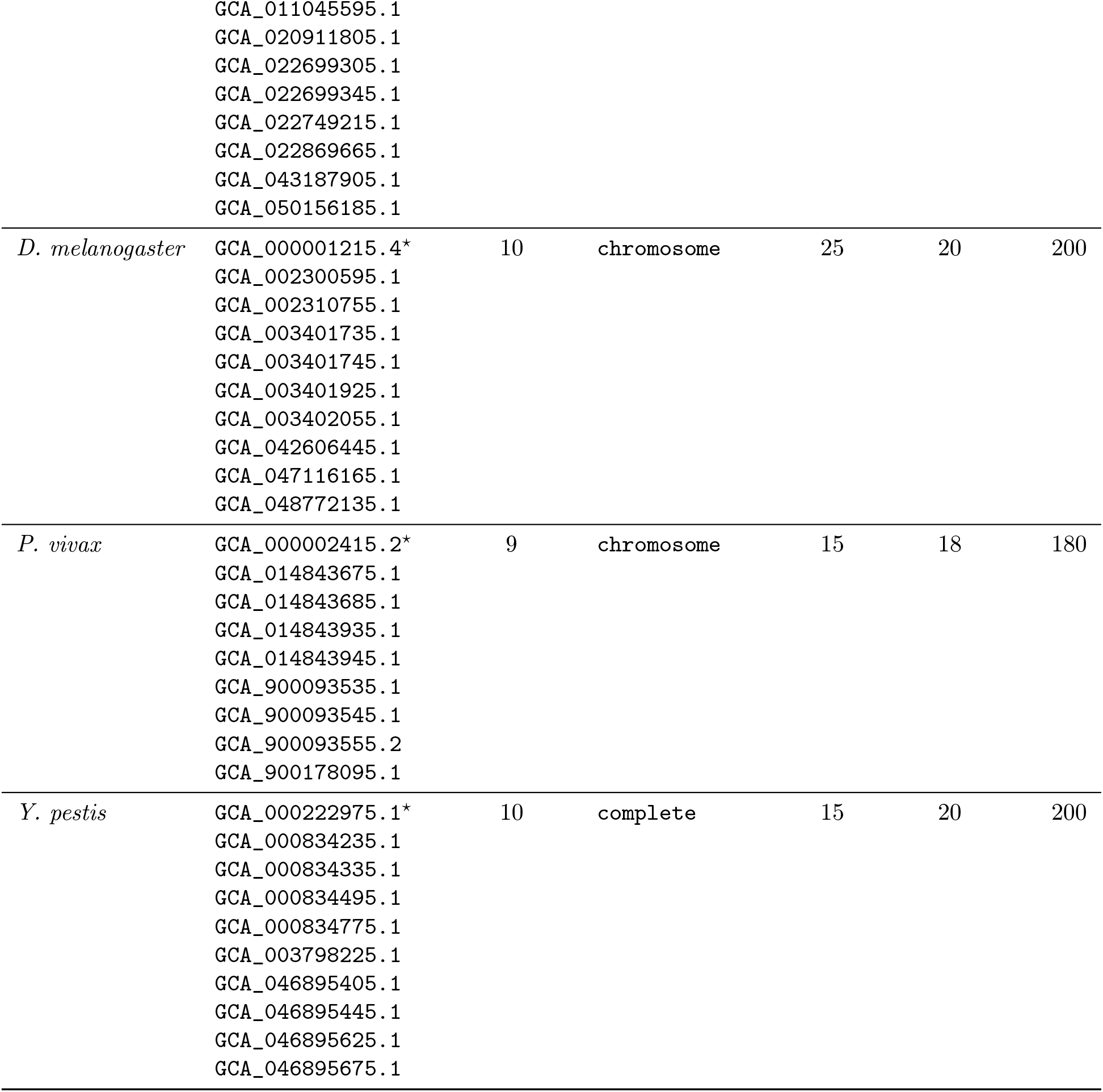
Genome datasets and segmentation settings used in Section 6.2. We list the accession numbers of the genomes used (`Accession.Version`), marking the reference used for Minigraph-Cactus with an asterisk (^⋆^). We summarize the number of genomes and the NCBI assembly level (Assembly lvl.). The last three columns describe parameters for SibeliaZ: the *k*-mer size used (*k* (SbZ)), the associated duplication filter parameters for the strong (*a* (SbZs)) and for the weak filtration (*a* (SbZw)) runs.

| Taxon | Accession.Version | #Genomes | Assembly lvl. | $k$ (SbZ) | $a$ (SbZs) | $a$ (SbZw) |
| --- | --- | --- | --- | --- | --- | --- |
| *A. thaliana* | GCA_000001735.2* | 10 | chromosome+ | 25 | 20 | 200 |
|  | GCA_020911765.2 |  |  |  |  |  |
|  | GCA_023115395.1 |  |  |  |  |  |
|  | GCA_028009825.2 |  |  |  |  |  |
|  | GCA_051624255.1 |  |  |  |  |  |
|  | GCA_051624265.1 |  |  |  |  |  |
|  | GCA_946405265.1 |  |  |  |  |  |
|  | GCA_946409395.1 |  |  |  |  |  |
|  | GCA_946409825.1 |  |  |  |  |  |
|  | GCA_965117575.1 |  |  |  |  |  |
| *C. carpio* | GCA_018340385.1* | 5 | chromosome+ | 25 | 10 | 100 |
|  | GCA_000951615.2 |  |  |  |  |  |
|  | GCA_027406505.1 |  |  |  |  |  |
|  | GCA_050881965.1 |  |  |  |  |  |
|  | GCA_051167465.1 |  |  |  |  |  |
| *S. cerevisiae* | GCA_000146045.2* | 10 | complete | 25 | 20 | 200 |
|  | GCA_001580425.1 |  |  |  |  |  |
|  | GCA_003086655.1 |  |  |  |  |  |
|  | GCA_004014915.1 |  |  |  |  |  |
|  | GCA_004328465.1 |  |  |  |  |  |
|  | GCA_021172205.1 |  |  |  |  |  |
|  | GCA_023508825.1 |  |  |  |  |  |
|  | GCA_024972935.1 |  |  |  |  |  |
|  | GCA_024972955.1 |  |  |  |  |  |
|  | GCA_903819175.2 |  |  |  |  |  |
| *E. coli* | GCA_000005845.2* | 10 | complete | 15 | 20 | 200 |
|  | GCA_000008865.2 |  |  |  |  |  |
|  | GCA_000010385.1 |  |  |  |  |  |
|  | GCA_000013265.1 |  |  |  |  |  |
|  | GCA_000019645.1 |  |  |  |  |  |
|  | GCA_000210475.1 |  |  |  |  |  |
|  | GCA_002853715.1 |  |  |  |  |  |
|  | GCA_003018455.1 |  |  |  |  |  |
|  | GCA_003697165.2 |  |  |  |  |  |
|  | GCA_048568945.1 |  |  |  |  |  |
| *K. pneumoniae* | GCA_000240185.2* | 10 | complete | 15 | 20 | 200 |
|  | GCA_006364295.1 |  |  |  |  |  |
|  | GCA_011045595.1 |  |  |  |  |  |
|  | GCA_020911805.1 |  |  |  |  |  |
|  | GCA_022699305.1 |  |  |  |  |  |
|  | GCA_022699345.1 |  |  |  |  |  |
|  | GCA_022749215.1 |  |  |  |  |  |
|  | GCA_022869665.1 |  |  |  |  |  |
|  | GCA_043187905.1 |  |  |  |  |  |
|  | GCA_050156185.1 |  |  |  |  |  |
| *D. melanogaster* | GCA_000001215.4* | 10 | chromosome | 25 | 20 | 200 |
|  | GCA_002300595.1 |  |  |  |  |  |
|  | GCA_002310755.1 |  |  |  |  |  |
|  | GCA_003401735.1 |  |  |  |  |  |
|  | GCA_003401745.1 |  |  |  |  |  |
|  | GCA_003401925.1 |  |  |  |  |  |
|  | GCA_003402055.1 |  |  |  |  |  |
|  | GCA_042606445.1 |  |  |  |  |  |
|  | GCA_047116165.1 |  |  |  |  |  |
|  | GCA_048772135.1 |  |  |  |  |  |
| *P. vivax* | GCA_000002415.2* | 9 | chromosome | 15 | 18 | 180 |
|  | GCA_014843675.1 |  |  |  |  |  |
|  | GCA_014843685.1 |  |  |  |  |  |
|  | GCA_014843935.1 |  |  |  |  |  |
|  | GCA_014843945.1 |  |  |  |  |  |
|  | GCA_900093535.1 |  |  |  |  |  |
|  | GCA_900093545.1 |  |  |  |  |  |
|  | GCA_900093555.2 |  |  |  |  |  |
|  | GCA_900178095.1 |  |  |  |  |  |
| *Y. pestis* | GCA_000222975.1* | 10 | complete | 15 | 20 | 200 |
|  | GCA_000834235.1 |  |  |  |  |  |
|  | GCA_000834335.1 |  |  |  |  |  |
|  | GCA_000834495.1 |  |  |  |  |  |
|  | GCA_000834775.1 |  |  |  |  |  |
|  | GCA_003798225.1 |  |  |  |  |  |
|  | GCA_046895405.1 |  |  |  |  |  |
|  | GCA_046895445.1 |  |  |  |  |  |
|  | GCA_046895625.1 |  |  |  |  |  |
|  | GCA_046895675.1 |  |  |  |  |  |

## 4 PAOs Effects on Multiplicities in the MBG

To see which pangenome structures a given MBG permits, we need to further characterize how many times a certain edge occurs in a pangenome. To that end it is helpful to classify the number of occurrences of an edge in a path.

Note in some cases *m*^*t*^*m*^*h*^ might be both a marker and an adjacency edge, leading to ambiguity which edge *m*^*t*^*m*^*h*^ refers to. In order to keep the notation simple, we assume such edges do not exist in the MBG. In cases where they do exist, one can easily replace the marker *m*^*t*^*m*^*h*^ by two markers 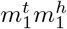 and 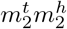, which are always neighboring with adjacency 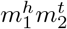 . (For Section 5.2, note that 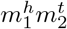 is not contested and is thus not affected by our algorithm cutting contested edges.)

If for a path or cycle *P* = *v*_1_ … *v*_n_ and an edge *uv* there is a *j*, 1 ≤ *j < n*, with {*v*_j_, *v*_j+1_} = {*u, v*} we say that *uv occurs* in *P* (at position *j*). If *P* is a cycle and {*v*_n_, *v*_1_} = {*u, v*}, we say that *uv* occurs at position *n*. We additionally define occ(*e, P*) as the total number of times an edge occurs in a path or cycle *P* .

For example, in path 1^*h*^2^*t*^2^*h*^2^*h*^2^*t*^1^*h*^1^*t*^3^*t*^, the edge 1^*h*^2^*t*^ occurs twice (at position 1 and at position 5), while the edge 2^*h*^2^*h*^ occurs once (at position 3).

For a pangenome ℙ and its MBG *G*, we write *m*_ℙ_(*uv*) for the total number of occurrences of the edge *uv* in ℙ. We call *m*_ℙ_ the *(edge) multiplicities*.

Given MBG *G* and multiplicities *m*, we write 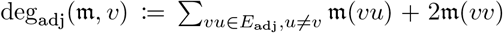 for the *adjacency degree* of a vertex *v* ≠∅, 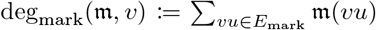 for the *marker degree* of a vertex *v* ≠∅, and deg_mark_(*m*, ∅) := 2*m*(∅∅), 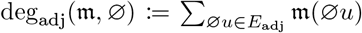 for telomere node ∅. When *m* is unambiguous, we omit it and simply write deg_adj_(*v*), deg_mark_(*v*). If edge multiplicities are derived from a pangenome, adjacency degree and marker degree coincide for any vertex *v*. We formalize this notion with the following definition.

### Definition 5.

*Given an MBG G* = (*V, E*_*adj*_ ∪ *E*_*mark*_) *and a multiplicity function m we call* (*G, m*) *an* alternating Eulerian graph *if* deg_*adj*_(*m, v*) = deg_*mark*_(*m, v*) *for all v* ∈ *V and m*(*e*) *>* 0 *for all e* ∈ *E*_*adj*_ ∪ *E*_*mark*_.

### 4.1 Alternating Eulerian graphs and pangenomes

We now show that an alternating cycle cover corresponding to the multiplicities exists if and only if *G* is alternating Eulerian.

#### Lemma 1.

*Given an MBG G and multiplicities m, there is an alternating cycle cover* ℂ *with m*(*e*) = ∑_C∈C_ *occ*(*e, C*) *if and only if* (*G, m*) *is an alternating Eulerian graph*.

In order to prove this lemma, an additional term is useful. Since we are interested in which cycle covers the graph permits, we distinguish paths that are compatible with the multiplicities from paths that are not compatible.

#### Definition 6.

*An alternating path or cycle P is called* valid *for multiplicities m if for each edge e that occurs in P holds m*(*e*) ≥ *occ*(*e, P*).

We will now show the equivalence between alternating Eulerian graphs and graphs that have an alternating cycle cover. In that regard, the following lemma is helpful.

#### Lemma 2.

*Given a valid path P in an alternating Eulerian graph, there is a valid path Q, such that* (*PQ*) *is a valid alternating cycle*.

*Proof*. We give an algorithm similar to the Hierholzer algorithm [24] to find *Q* in Algorithm 1.

Multiplicities *m*′ ensure that the path is valid while the case distinction ensures that *Q* is alternating.

Thus the only potential problem is whether edge *vx* exists. Let *uv* be an adjacency edge. Say *PQ* contains a total number of *l* adjacency edges at *v*. Then, either the first edge in *P* is a marker edge starting at *v* or *PQ* contains *l* − 1 occurrences of marker edges adjacent to *v*. In the former case, *PQ* is already an alternating cycle, so the loop would not have been started. In the latter case, observe that due to the graph being alternating Eulerian, deg_mark_(*v*) = deg_adj_(*v*) ≥ *l* and therefore a marker edge *vx* with *m*′[*vx*] *>* 0 must exist. The case for a marker edge *uv* is analogous.

□

#### Algorithm 1

Algorithm to complete a valid path to a valid cycle.

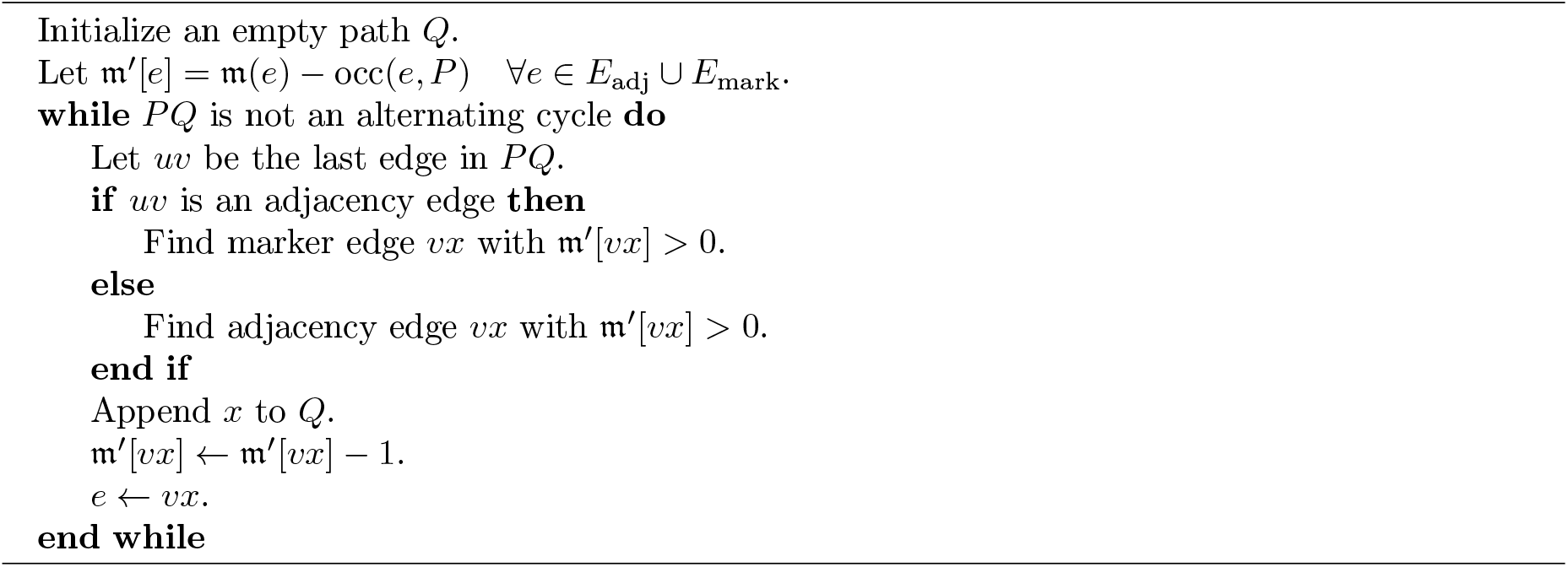

We can now prove Lemma 1:

*Proof*. Assume there is an alternating cycle cover. In each cycle, if a vertex *v* is preceded by an adjacency edge, it is succeeded by a marker edge and vice versa. Thus, deg_adj_(*v*) = deg_mark_(*v*).

Using Lemma 2, creating an alternating cycle cover is easy. Take an edge *uv* with *m*(*uv*) *>* 0. Then there must be a path *Q*, such that *C* := (*uvQ*) is a valid alternating cycle. Then adjusting the multiplicities *m*′(*e*) := *m*(*e*) −occ(*e, C*) and removing edges *e* with *m*′(*e*) = 0, yields another alternating Eulerian graph. Applying this procedure iteratively yields the cycle cover.

### 4.2 Diminishing paths

Due to Lemma 1, if we want to find a pangenome that has a particular MBG *G*, we only need to find a multiplicity function *m*, such that (*G, m*) is alternating Eulerian. Particularly for applying GAOs most effectively, we want to have a pangenome where certain edges are as rare as possible. Since we start with an alternating Eulerian graph, we only need to find a modified multiplicity function which is smaller for the desired edges.

As an example of how we reduce multiplicities, regard the following path: 1^*t*^1^*h*^2^*t*^2^*h*^ where *m*(1^*t*^1^*h*^) = 3, *m*(1^*h*^2^*t*^) = 2 and *m*(2^*t*^2^*h*^) = 4. Rewriting the multiplicities to *m*′(1^*t*^1^*h*^) := *m*(1^*t*^1^*h*^) −1 = 2, *m*′(1^*h*^2^*t*^) := *m*(1^*h*^2^*t*^) −1 = 1 and *m*′(2^*t*^2^*h*^) := *m*(2^*t*^2^*h*^) −1 = 3 ensures that the condition of Definition 5 is still met for *m*′ at vertices 1^*h*^ and 2^*t*^, but not at the ends of the path, 1^*t*^ and 2^*h*^. If we can find a cycle with this property, we can thus reduce the multiplicities while the graph stays alternating Eulerian. Generally, such paths can be defined as follows.

#### Definition 7.

*An alternating path or cycle P in MBG G with multiplicities m is called* diminishing *if for each edge e* ∈ *E*_*adj*_ ∪ *E*_*mark*_ *holds that occ*(*e, P*) *< m*(*e*). *We call* 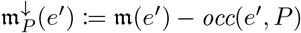 *the* diminution *of P* .

The following lemma generalizes the notion of how to reduce edge multiplicities while keeping the graph alternating Eulerian.

#### Lemma 3.

*Given a diminishing path P* = *v*_1_ … *v*_n_ *in an alternating Eulerian graph* (*G, m*) *in* 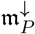 *holds* 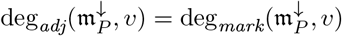 *for v* ∈ *V if v* ∉ {*v*_1_, *v*_n_}.

*Proof*. Let *v* ∉ {*v*_1_, *v*_n_} and *L*_v_ := {2 ≤ *i* ≤ *n* − 1 : *v*_i_ = *v*}. Then for *i* ∈ *L*_v_, exactly one of *v*_i−1_*v*_i_ and *v*_i_*v*_i+1_ is an adjacency edge and exactly one is a marker edge. Therefore, 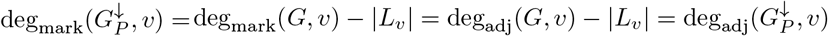.

□

We can immediately derive the following corollary.

#### Corollary 1.

*Given a diminishing cycle C* = (*v*_1_ … *v*_n_) *in an alternating Eulerian graph* (*G, m*), 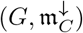 *is alternating Eulerian*.

Our goal is thus to reduce multiplicities using diminishing cycles. In this context, it is useful to recognize when a path already contains a diminishing cycle. The following concept helps to detect this situation.

#### Definition 8.

*An alternating path or cycle P is* reducible *if it is of the form P* = *s*_0_*m*^*t*^*m*^*h*^*s*_1_*m*^*t*^*m*^*h*^*s*_2_ *or P* = *s*_0_*m*^*h*^*m*^*t*^*s*_1_*m*^*h*^*m*^*t*^*s*_2_ *for some paths s*_0_, *s*_1_, *s*_2_ *and some marker m. Otherwise, P is* irreducible.

For a diminishing reducible path, such as *P* = *s*_0_*m*^*t*^*m*^*h*^*s*_1_*m*^*t*^*m*^*h*^*s*_2_, we can then instead consider the diminishing cycle (*m*^*t*^*m*^*h*^*s*_1_). Note that this also holds for paths of the form *s*_0_∅∅*s*_1_∅∅*s*_2_.

### 4.3 A complete set of Path Altering Operations

We now show that there is a set of operations, which is a complete set of PAOs and describes meaningful biological interactions. In fact, we do so by extending the PAOs already described in Section 3.1. We only add a direct way by which genomes interact and an operation modeling chromosomal recombination.

While there are many ways in which genomes can interact, the most direct one is by transferring chromosomal material.

#### Definition 9

(Chromosome Transfer). *Given a pangenome* ℙ *that includes genomes G*_1_, *G*_2_ *and a multiset of chromosomes C* ⊆ *G*_1_, *a chromosome transfer between G*_1_ *and G*_2_ *creates* ℙ′ = ℙ ⊖ {*G*_1_, *G*_2_} ⊕ 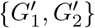 *where* 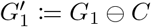 *and* 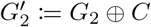.

If multiple copies of the same chromosome exist in the same genome, new chromosome structures can arise by recombinations, such as shown in Fig. 4. These homologous recombinations are one mechanism of how rearrangements occur [40].

**Figure 4:**
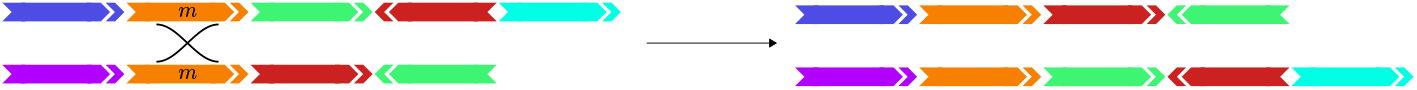
A homologous recombination may create new chromosome structures not previously present in the pangenome.

We can define a homologous recombination in our context roughly as taking two (sub-)strings *SmT* and *UmV, m* ∈ Σ, of one or two chromosomes and transforming them into *SmV* and *UmT* . Depending on the connectivity of these substrings, this then has different effects. More formally:

#### Definition 10.

*A* homologous recombination *transforms chromosomes of the same genome in one of the following ways*

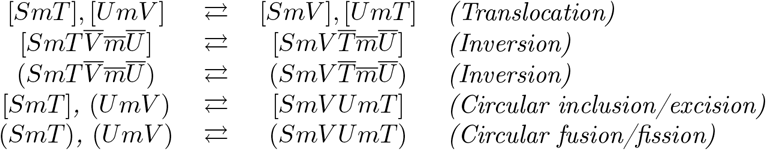

*where m* ∈ Σ *and S, T, U*, 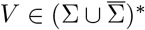.

Note that a homologous recombination is a PAO, because while the chromosome connectivity is transformed, the adjacencies (particularly at marker *m*) remain the same and therefore the graph remains unchanged.

Transfer and recombination in conjunction with chromosome (de-) multiplication, coalescence and divergence are then a complete set of PAOs.

#### Theorem 1.

*The set of operations Ñ, which consists of (de-) multiply, homologous recombination, coalescence, divergence and chromosome transfer operations is a complete set of PAOs*.

The proof for this theorem rests on the fact that each operation has an inverse in *Ñ*. It thus suffices to show that any pangenome ℙ_a_ can be transformed into 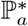 where 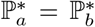 if and only if *G*(ℙ_a_) = *G*(ℙ_b_).

We begin by deriving a few helpful lemmas.

#### Lemma 4.

*There is a finite number of irreducible cycles in a multiple breakpoint graph*.

*Proof*. Each irreducible cycle can contain any marker at most twice, as *m* and 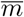. Then for any irreducible cycle we can assign a unique string 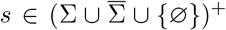 with |*s*| *≤* 2 |Σ| + 2. Since the number of such strings is finite (finite length and finite alphabet), the number of irreducible cycles is finite as well.

□

We further show that a cycle can always be decomposed into a set of irreducible cycles.

#### Lemma 5.

*Any cycle C can be transformed into a set I*(*C*) *of irreducible cycles by homologous recombinations*.

*Proof*. If *C* is irreducible, the set is *I*(*C*) = {*C}*. If *C* = (*S*_1_*m*^*t*^*m*^*h*^*S*_2_*m*^*t*^*m*^*h*^) for some marker *m* and paths *S*_1_ and *S*_2_, we perform the homologous recombination (*S*_1_*m*^*t*^*m*^*h*^*S*_2_*m*^*t*^*m*^*h*^) → *C*_1_ := (*S*_1_*m*^*t*^*m*^*h*^), *C*_2_ := (*S*_2_*m*^*t*^*m*^*h*^), recursively apply the procedure to *C*_1_ and *C*_2_, and finally set *I*(*C*) = *I*(*C*_1_) ∪ *I*(*C*_2_).

□

Let *I*^*^ be the set of all irreducible cycles in the MBG of ℙ. We then define ℙ* := {*I*^*^} as the pangenome consisting of a single genome (*I*^*^). Note that 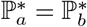 if and only if *G*(ℙ_a_) = *G*(ℙ_b_).

#### Lemma 6.

*Any pangenome* ℙ *can be transformed into* ℙ^+^ = {*I*}, *a pangenome consisting of a single genome I with only irreducible cycles by operations in Ñ*.

*Proof*. We first show how to unify all genomes of the pangenome into a single genome. Consider two genomes *G*_1_, *G*_2_. Let *A*_1_ := *G*_2_ ⊖ *G*_1_ and *A*_2_ := *G*_1_ ⊖ *G*_2_. We can then create 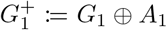 by duplicating the chromosomes from *A*_1_ in *G*_2_ and transferring them to *G*_1_. Analogously, we can create 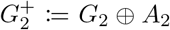. Then, 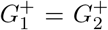. We then unify 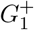 and 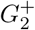 via coalescence. One can iterate this procedure until there is a single genome *G*^+^, which can then be transformed into *I* containing only irreducible cycles by Lemma 5.

□

#### Lemma 7.

*Given a genome consisting of irreducible cycles I and an irreducible cycle C in G*({*I*}) *with C* ∉ *I, there is a series of duplications and homologous recombinations transforming I into a set of irreducible cycles I*′ *with C* ∈ *I*′ *and I* ⊂ *I*′.

*Proof*. Let *C* = (*m*_1_*m*_2_ … *m*_n_) where each *m*_i_ is some marker in forward or backward orientation. Since all adjacencies of *C* are from the MBG of {*I*}, there are cycles *C*_1_, …, *C*_n_ ∈ *I* with *C*_1_ = (*m*_1_*m*_2_*S*_1_), *C*_2_ = (*m*_2_*m*_3_*S*_2_), …, *C*_n_ = (*m*_n_*m*_1_*S*_n_) for some paths *S*_1_ … *S*_n_. We duplicate all cycles *C*_1_, …, *C*_n_. Then, starting with *C*_1_, *C*_2_, we perform homologous recombinations of the following type:

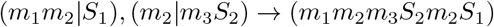

until we merge all cycles into a single cycle with the following structure:

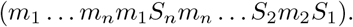

We then perform the homologous recombination

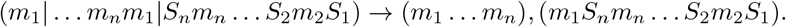

The remaining cycle (*m*_1_*S*_n_*m*_n_ … *S*_2_*m*_2_*S*_1_) can then be transformed into irreducible cycles (see Lemma 5). Now *I*′ contains only irreducible cycles, *C* = (*m*_1_ … *m*_n_) ∈ *I*′, and since all cycles we operated on had a duplicate, we also have *I* ⊂ *I*′.

Using Lemmas 6 and 7, we can derive the following corollary.

#### Corollary 2.

*Any pangenome* ℙ *can be transformed into* ℙ* *by operations in Ñ*.

Since each operation has an inverse in *Ñ*, this proves Theorem 1. Note that this is by no means the shortest or most realistic scenario. Optimizing for the minimum number of PAOs to transform one pangenome into another one may be subject to future research.

Also note that we only discussed a small subset of PAOs here. There are many more possible PAOs. Some rearrangement operations which have been studied recently, such as flanked transpositions [50] or flanked block interchanges [30], are in fact PAOs, although none of them form a complete set by themselves. Nonetheless, this means the complete set of PAOs *Ñ* discussed here is not the only such set. However, since PAOs are used without a cost in Problem 1, the concrete set of PAOs does not change the CARP measure or CARP simplification as long as the set is complete.

## 5 Solution under the SCJ model

The *Single Cut or Join (SCJ)* model, established by Feijão and Meidanis [18], is a simple rearrangement model closely related to the breakpoint distance. The model allows to cut a chromosome in one place (single cut) or to join two ends from one or two linear chromosomes together (join). These two operations have analogues in biology, namely double strand breaks and non-homologous end joining. However, care needs to be taken in interpreting genomes reconstructed by SCJ since events with breakpoint reuse are not reflected in SCJ scenarios.

Nonetheless, it is a useful model as many otherwise NP-hard problems are tractable under SCJ. However, reconstruction without a tree (large parsimony problem), remains NP-hard [18]. One might thus expect that CARP is NP-hard as well under SCJ. However, we show here that the problem has a surprisingly simple linear time solution.

Formally, an SCJ is either a *Single Cut* or a *Single Join*. A Single Cut transforms a circular chromosome (*S*) into a linear chromosome [*S*] or a linear chromosome [*ST*] into two linear chromosomes [*S*] and [*T*]. A *Single Join* transforms a linear chromosome [*S*] into a circular chromosome (*S*) or two linear chromosomes [*S*] and [*T*] into one linear chromosome [*ST*]. *S* and *T* here represent arbitrary strings of oriented markers.

### 5.1 Lower bound

In the MBG, cutting between two markers *m, n*, i.e. [*SmnT*] → [*Sm*], [*nT*], creates the telomeric adjacencies *m*^*h*^∅, *n*^*t*^∅ and removes the adjacency edge *m*^*h*^*n*^*t*^ if [*SmnT*] was the only chromosome with that adjacency. Conversely, the join [*Sm*], [*nT*] → [*SmnT*] creates the adjacency edge *m*^*h*^*n*^*t*^ and removes *m*^*h*^∅, *n*^*t*^∅ if [*Sm*], [*nT*] are the only linear chromosomes with these (telomeric) adjacencies. The effects for other combinations of orientations and chromosome types are analogous. Note that if *uv* and *u*′*v*′ are joined to *uu*′, then *v* = *v*′ = ∅.

From this follows that, to arrive at a simple pangenome, we need to expend at least one SCJ operation per non-telomeric adjacency that involves a branching node. We call these adjacencies *contested*. We give an example graph where contested edges are marked in Fig. 5.

**Figure 5:**
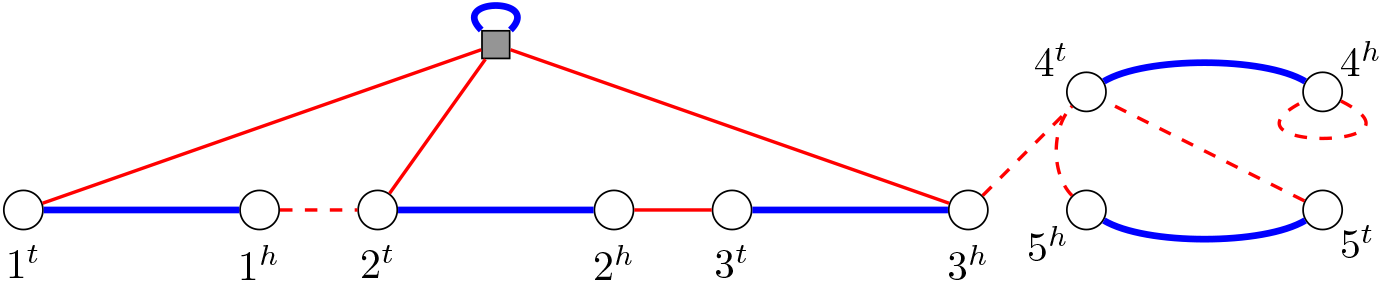
MBG for the pangenome 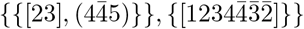 with contested edges shown dashed. Blue edges are marker edges, red edges are adjacency edges, and the quadratic grey node is the telomere ∅.

#### Lemma 8.

*Transforming a pangenome* ℙ *with MBG G*(ℙ) = (*V, E*_*adj*_∪*E*_*mark*_) *into a simple pangenome* A *requires at least* |*E*_*cnt*_| *SCJ operations where E*_*cnt*_ := {*uv* ∈ *E*_*adj*_ | *u* = *v*} ∪ {*uv* ∈ *E*_*adj*_ | *u, v*≠ ∅, ∃*uv*′ ∈ *E*_*adj*_ *with v*′≠ *v*}. *We call E*_*cnt*_ *the set of* contested adjacencies.

*Proof*. We prove this lemma using an injection *f* between edges in *E*_cnt_ and SCJ operations in the CARP scenario. Consider an edge *uv* ∈ *E*_cnt_. We distinguish two cases.

I. Let *uv* not be an adjacency in the simple MBG of A. Then there must be a cut *c*_uv_ removing *uv* from the graph before A is reached, i.e. 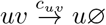, *v*∅. We thus map *uv* to *c*_uv_, i.e. *f* (*uv*) = *c*_*uv*_ .
II. Let *uv* be an adjacency in the simple MBG of A. Note that then *u* ≠ *v*, otherwise the final MBG would not be simple. Since *uv* is contested, there is another adjacency edge at *u* or at *v*. Let w.l.o.g. *uv*′ ∈ *E*_adj_ with *v*′≠ *v*. In the CARP scenario, we may then have a series of alternating cuts and joins exchanging *v*′ for other nodes: 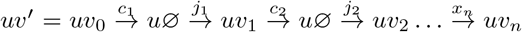 . The MBG of A, which contains, is simple only if *v*_n_ = *v* ≠ . Therefore the last operation in the series must be a join, *x*_n_ = *j*_n_. We thus map *uv* to *j*_n_, i.e. *f* (*uv*) = *j*_n_.

□

We thus need to expend at least one SCJ operation per contested edge. In the following we show that this lower bound is tight.

### 5.2 Upper Bound

In principle, the bound in Lemma 8 can be reached by applying a cut *uv* → *u*∅, *v*∅ to each contested edge *uv*. However note that if *uv* occurs more than once in the pangenome, we would need multiple cuts to remove it from the pangenome. Therefore we use the complete set of PAOs to ensure there is a contested edge occurring exactly once, arriving at the following central theorem.

#### Theorem 2.

*The CARP measure under the SCJ model for a pangenome* P *with MBG G*(P) = (*V, E*_*adj*_∪ *E*_*mark*_) *is* |*E*_*cnt*_|. *One possible SCJ-CARP simplification* A *is defined by the adjacencies E*_A_ := *E*_*adj*_ \ E_*cnt*_

Using the concepts introduced in Section 4.2, we now show that for any graph with contested edges, there is a pangenome that contains some *e* ∈ *E*_cnt_ with *m*(*e*) = 1. The first step is to show that there are diminishing cycles or edges with multiplicity 1. To prove this, we derive a number of useful facts, which we summarize in the following lemma.

#### Lemma 9.

*Let P* = *v*_1_ … *v*_n_ *be a diminishing path in an alternating Eulerian graph* (*G, m*), *such that*

- (*G, m*) *does not contain any diminishing cycles*
- *P cannot be extended to the right, i*.*e*., *there is no path P* ′ = *Pu that is diminishing*.

*Let U be the set of vertices with u* ∈ *U* ⇐⇒ *Pu is an alternating path, and let U*_≥2_ ⊆ *U be the set of vertices u* ∈ *U for which m*(*v*_n_*u*) ≥ 2. *Then all of the following hold:*

i. ∀*u* ∈ *U*_≥2_, *v*_n_*u occurs in P at some position i and only in reverse order, i*.*e. v*_i_ = *u, v*_i+1_ = *v*_n_.
ii. *v*_n_ ∈ *U*_≥2_,
iii. |*U*_≥2_| ≤ 1,
iv. ∀*u* ∈ *U*_≥2_, *v*_n_*u occurs exactly once in P*,
v. ∀*u* ∈ *U*_≥2_ *holds m*(*v*_n_*u*) = 2,
vi. ∃*u* ∈ *U with m*(*v*_n_*u*) = 1.

*Proof*. In the following note that the graph not containing diminishing cycles implies that *P* is irreducible.

i. If there is any *u* ∈ *U*_≥2_, such that *v*_n_*u* does not occur in *P*, then *v*_1_ … *v*_n_*u* is a diminishing path, a contradiction. Therefore for all *u* ∈ *U*_≥2_, *v*_n_*u* occurs in *P* . If there is any *u* ∈ *U*_≥2_, such that there is 1 ≤ *j < n* with *v*_j_ = *v*_n_ and *v*_j+1_ = *u*, then (*v*_j+1_ … *v*_n_) is a diminishing cycle, a contradiction. Thus for each *u* ∈ *U v*_n_*u* occurs in *P* and only in reverse order. □_(i)_
ii. Assume *v*_n_ ∈ *U*_≥2_. Then, using (i), there is some *i* with *v*_i_*v*_i+1_ = *v*_n_*v*_n_ and thus (*v*_i+1_ … *v*_n_) is a diminishing cycle, a contradiction. □_(ii)_
iii. Let there be *u, u*′ ∈ *U*_≥2_ with *u* = *u*′. Then, *v*_n_*u* and *v*_n_*u*′ must be adjacency edges, because there is exactly one marker edge incident to each vertex. Using (i), both occur in reverse in *P* . W.l.o.g. let *uv*_n_ occur before *u*′*v*_n_ in *P* . Then we have *P* = … *uv*_n_ … *u*′*v*_n_ … *v*_n_. Since *v*_n_*u*′ is an adjacency edge, *v*_n−1_*v*_n_ must be a marker edge. Since each vertex is adjacent to exactly one marker edge, we have *P* = … *uv*_n_*v*_n−1_ … *u*′*v*_n_*v*_n−1_ … *v*_n_, meaning that *P* is reducible, a contradiction. □_(iii)_
iv. For *u* ∈ *U*_≥2_, let *uv*_n_ occur twice in *P* . As before, we then have *P* = … *uv*_n_*x* … *uv*_n_*y* … *v*_n_. Either *uv*_n_ is a marker edge or *v*_n_*x* is a marker edge, in which case *v*_n_*x* = *v*_n_*y*. In both cases *P* is reducible, a contradiction. □_(iv)_
v. Because *u* ∈ *U*_≥2_ occurs only once in *P*, if *m*(*v*_n_*u*) *>* 2, *Pu* would be a diminishing path. □_(v)_
vi. Using (i) and (ii) we know that for *u* ∈ *U*_≥2_, either (a) *P* = … *uv*_n_*v*_j+2_ … *v*_n−1_*v*_n_ for some *j*, or (b) *P* = … *uv*_n_*v*_n_ and with (iii) that *U*_≥2_ = {*u*}.

Regard the adjacency and marker degree of *v*_n_. In (a), *v*_n_*v*_j+2_ and *v*_n−1_*v*_n_ have the same type. We then distinguish the cases:

I. *v*_j+2_ ≠ *v*_n−1_. Then *v*_n−1_*v*_n_ and *v*_n_*v*_j+2_ are adjacency edges. Since *P* is diminishing *m*(*v*_n−1_*v*_n_) *>* 1 and *m*(*v*_n_*v*_j+2_) *>* 1. Thus deg_adj_(*v*_n_) ≥ *m*(*v*_n−1_*v*_n_) + *m*(*v*_n_*v*_j+2_) ≥ 4
II. *v*_j+2_ = *v*_n−1_. Then because *P* is diminishing *m*(*v*_n−1_*v*_n_) *>* 2. Thus deg_adj_(*v*_n_) = deg_mark_(*v*_n_) ≥ *m*(*v*_n−1_*v*_n_) ≥ 3.

For (b), we have deg_adj_(*v*_n_) = deg_mark_(*v*_n_) ≥ 2*m*(*v*_n_*v*_n_) ≥ 4.

In all cases (a(I), a(II), b), we have deg_adj_(*v*_n_) = deg_mark_(*v*_n_) *>* 2, and since *U*_≥2_ = {*u*} with *m*(*v*_n_*u*) = 2 (see (v)), there must be at least one other edge *v*_n_*u*′ ∉ *U*_≥2_ that has therefore *m*(*v*_n_*u*′) = 1. If |*U*_≥2_| = 0, all edges *v*_n_*u* with *u* ∈ *U* have *m*(*v*_n_*u*) = 1. □_(vi)_

□

The rough idea for the algorithm cutting contested edges (see also Algorithm 2) is thus as follows. As long as there are contested edges of multiplicity 1, we cut them. If we cannot find such an edge any more, we find diminishing cycles, which we then use to reduce the multiplicities in the graph until there is an edge of multiplicity 1, so we can repeat the process. However, it may be the case that none of the edges with multiplicity 1 is contested. If that is the case, a slightly more complex procedure needs to be employed.

We begin by showing that a contested edge *uv* with some helpful properties always exists.

#### Lemma 10.

*For any alternating Eulerian MBG* (*G, m*) *where m*(*e*) *>* 1 ∀*e* ∈ *E*_*cnt*_, |*E*_*cnt*_| ≠ 0 *and* (*G, m*) *has no diminishing cycles, there is an edge v*∅ ∈ *E*_*adj*_ *with m*(*v*∅) = 1, *such that uv* ∈ *E*_*adj*_, *u* ≠ *v with m*(*uv*) = 2 *and u*, ∅ *the only vertices adjacent to v*.

*Proof*. Regard an (irreducible) diminishing path *P* = *v*_1_ … *v*_n_ that cannot be extended in either direction, i.e., there is no path *av*_1_ … *v*_n_ or *v*_1_ … *v*_n_*b* that is diminishing. Then according to Lemma 9 (vi), there are edges *xv*_1_, *v*_n_*y* with multiplicity 1. (Note that this may also be the same edge.)

Regard *v*_n_*y*. Since *m*(*v*_n−1_*v*_n_) *>* 1 and *m*(*v*_n_*y*) = 1, there must be another edge *v*_n_*z* or *y* = *v*_n_. Analogously, there is *wv*_1_ or *x* = *v*_1_.

Therefore *xv*_1_, *v*_n_*y* are either adjacency edges or the telomere marker edge ∅∅. Assume now that *xv*_1_ = ∅∅. This implies deg_mark_(∅) = 2 and, since *m*(*v*_1_*v*_2_) ≥ 2, this means that *v*_1_*v*_2_ = ∅*v*_2_ is the only adjacency edge incident to ∅ with *m*(*v*_1_*v*_2_) = 2. However, then {*v*_n_, *y*} ≠ {*v*_2_, ∅}, meaning that *v*_n_*y* is contested, a contradiction. We may thus assume that both *xv*_1_ and *v*_n_*y* are adjacency edges and therefore *v*_1_*v*_2_ and *v*_n−1_*v*_n_ are marker edges.

If both *v*_1_ = *v*_n_ = ∅, since *v*_1_*v*_2_ and *v*_n−1_*v*_n_ are marker edges, *P* = ∅∅*v*_3_ … ∅∅ is reducible, a contradiction.

Let thus w.l.o.g. *v*_n_ ≠ ∅. Then *y* = ∅, otherwise *v*_n_*y* would be a contested edge with *m*(*v*_n_*y*) = 1. This means that *v*_n_*z* is a contested edge with *m*(*v*_n_*z*) *>* 1. According to Lemma 9 (iii) and (v) *v*_n_*z* has *m*(*v*_n_*z*) = 2 and there are no other edges with multiplicity 2 or higher adjacent to *v*_n_. Since any other edge *v*_n_*w* with *w* ≠ ∅ and *w* ≠ *z* would be contested and have *m*(*v*_n_*w*) = 1, such an edge does not exist. Additionally, due to Lemma 9 (ii), *u* ≠ *v*.

There is then a diminishing path that contains *uv* and that contains *v* only once.

#### Lemma 11.

*For any alternating Eulerian MBG* (*G, m*) *where m*(*e*) *>* 1 ∀*e* ∈ *E*_*cnt*_, |*E*_*cnt*_| ≠ 0 *and* (*G, m*) *has no diminishing cycles, for a pair of vertices u, v as described in Lemma 10, there is a diminishing path vuu*_1_ … *u*_m_ *that cannot be extended to the right, such that v does not occur in u*_1_ … *u*_m_.

*Proof*. Observe that the single edge *vu* is already a diminishing path. Therefore *vuu*_1_ … *u*_m_ exists.

We now show by contradiction that *v* does not occur in *u*_1_ … *u*_m_. Let *v*′ be the other extremity of the marker of *v*. We distinguish two cases:

I. *v* occurs before *v*′ in *u*_1_ … *u*_m_ at position *k*, that is we have *u*_1_ … *u*_k−1_*v*, where *u*_k−1_*v* is an adjacency edge. Then consider *u*_k−1_. According to Lemma 10, *u*_k−1_ = ∅ or *u*_k−1_ = *u*. In both cases, *vuu*_1_ … *u*_k−1_*v* is not diminishing, a contradiction.
II. *v*′ occurs before *v* in *u*_1_ … *u*_m_ at position *l*. Then we have *u*_1_ … *u*_l−1_*v*′*v*. However, then (*v*′*vuu*_1_ … *u*_l−1_) is a diminishing cycle, a contradiction.

□

We are now ready to construct the multiplicity function *m*′, in which *uv* has multiplicity 1.

#### Lemma 12.

*For any alternating Eulerian MBG* (*G, m*) *where m*(*e*) *>* 1 ∀*e* ∈ *E*_*cnt*_, |*E*_*cnt*_| ≠ 0 *and* (*G, m*) *has no diminishing cycles, there is m*′, *such that* (*G, m*′) *is alternating Eulerian and* ∃*uv* ∈ *E*_*cnt*_ *with m*′(*uv*) = 1.

*Proof*. Let *u, v* be vertices as described in Lemmas 10 and 11, i.e., *v*∅ ∈ *E*_adj_ and *m*(*vu*) = 2, and let *P* = *vuu*_1_ … *u*_m_ be the diminishing path that cannot be extended further to the right. Then, according to Lemma 9 (vi), there is an edge *u*_m_*x* with *m*(*u*_m_*x*) = 1. We distinguish two cases, visualized in Fig. 6.

**Figure 6:**
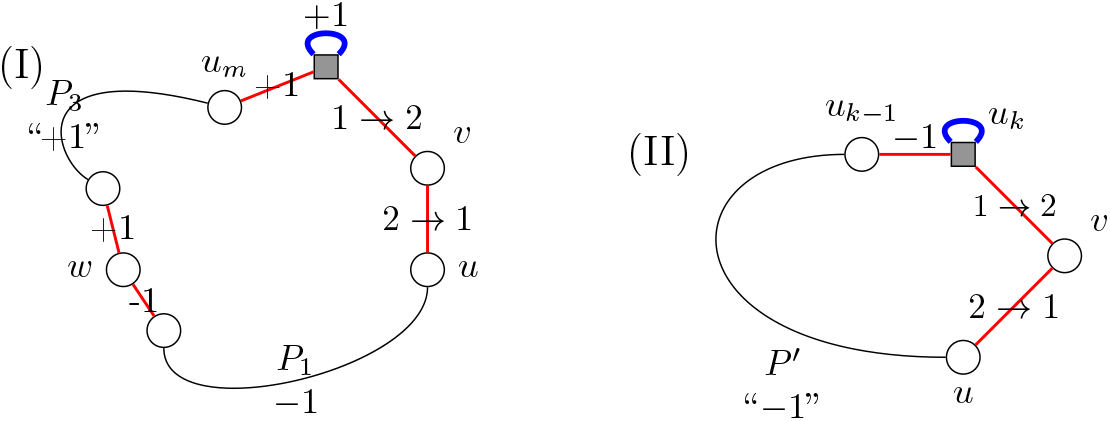
Case distinction in proof of Lemma 12. Black lines represent arbitrary alternating cycles that do not share edges or vertices. Note that the edges shown explicitly do not occur in the black paths.

(I) ∅ does not occur in *P* . Then *x* = ∅. Then there exists another edge *u*_m_*z*, which is contested. It therefore has multiplicity *m*(*u*_m_*z*) = 2 and occurs once in *P*_vu_ (see Lemma 9 (iv),(v)). Thus at least one vertex (*u*_m_) occurs twice in *P* . Let *w* be the leftmost vertex that occurs more than once in *P* . Let *w*′ be the other extremity of the marker of *w*. Then *P* is of the form *P* = *P*_1_*ww*′*P*_2_*w*′*wP*_3_ with *P*_1_ containing only vertices that occur once in *P* and *P*_3_ not containing further occurrences of *w*. Note that because of this, *P*_1_ and *P*_3_ do not share any vertices. We then set

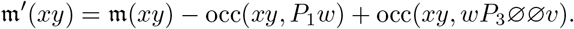

Note that there are only two vertices at which the condition for an alternating Eulerian path might not hold, namely *v* and *w*. Regard the adjacency degree of *v*.

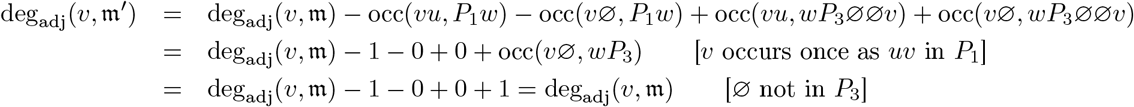

Let *v*′ be the other extremity of the marker of *v*. Note that because *v* occurs only once in *P* at the beginning in an adjacency edge, *vv*′ does not occur in *P* .

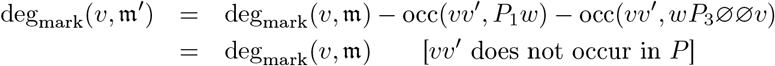

We thus see that at *v* the degree is unchanged and the condition for an alternating Eulerian graph still holds.

Regard the adjacency and marker degree of *w*. By definition *w* does not appear in *P*_1_, *P*_3_. Therefore, its marker degree does not change and for the adjacency degree holds

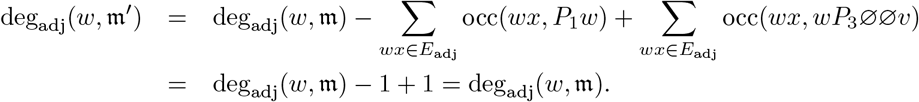

Thus, (*G, m*′) is alternating Eulerian. Observe that the contested edge *uv* now has multiplicity *m*′(*uv*) = *m*(*uv*) − occ(*uv, P*_1_*w*) + occ(*uv, P*_3_∅∅*v*) = 2 − 1 + 0 = 1.

(II) ∅ occurs in *P* . Then let *u*_k_ = ∅ be the first occurrence of ∅ in *P* and *P* ′ = *vuu*_1_ … *u*_k−1_∅. Note that *u*_k−1_≠ *v* because *m*(*u*_k−1_∅) *>* 1 (*P* is diminishing). We then set

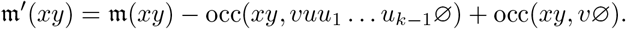

Again note that the adjacency and marker degrees of all nodes except *v* and ∅ under *m*′ are in agreement. The marker degrees of *v* and ∅ do not change as the markers do not occur in *vuu*_1_ … *u*_k−1_∅ or *v*∅.

Regard the adjacency degree of *v*.

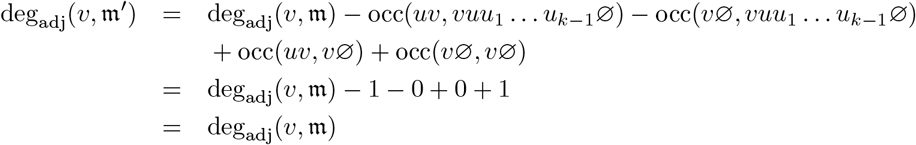

Regard the adjacency degree of ∅.

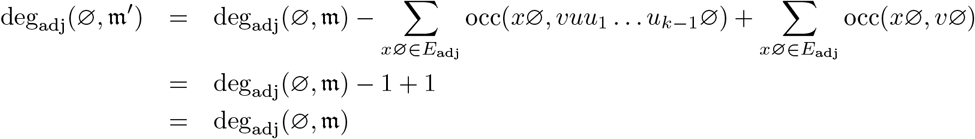

Therefore (*G, m*′) is alternating Eulerian. The multiplicity of *uv* is

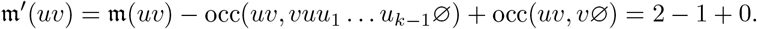

□

Using Lemma 1, Corollary 1 and Lemma 12 we can thus ensure that as long as there are contested adjacencies, there is an edge *e* ∈ *E*_cnt_ with *m*(*e*) = 1. The lower bound in Lemma 8 can thus be reached by cutting all edges in *E*_cnt_, proving Theorem 2.

The following algorithm shows how the GAO scenario for SCJ-CARP can be reconstructed. Future research might examine how to reconstruct the shortest PAO scenario. The results from this research could then be used here (see Line 19) to reconstruct both the PAO and GAO sequences. To simply reconstruct a simplification and the number of SCJs in the scenario, Algorithm 2 is not necessary; it suffices to compute *E*_cnt_ (see Theorem 2).

#### Algorithm 2

SCJ-CARP scenario reconstruction

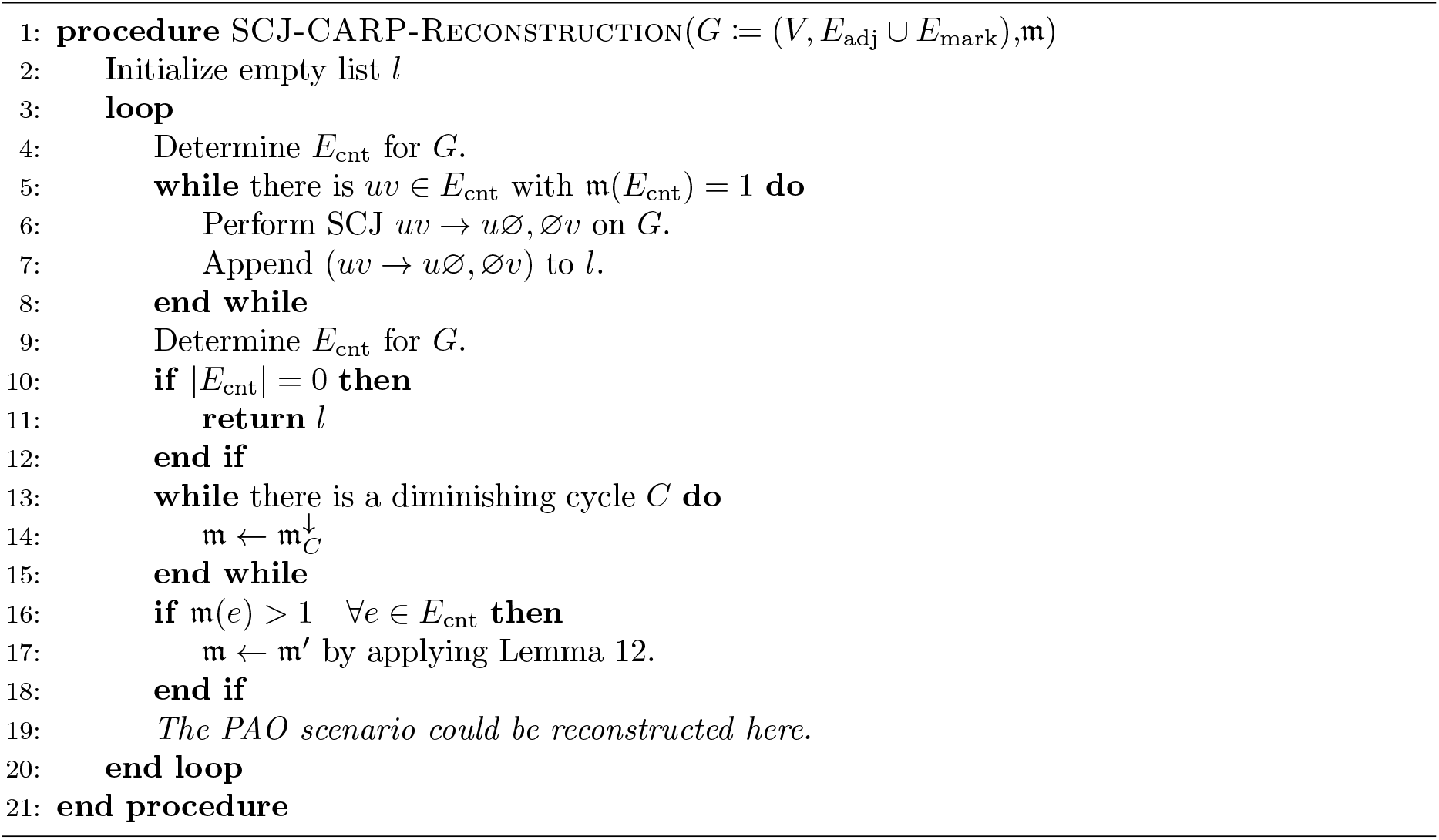

CARP under SCJ is thus solvable in time linear with respect to the graph size. Notably, the CARP measure corresponds directly to an easy to count graph metric, namely the edges connecting to branching nodes *E*_cnt_. Indeed, this mirrors the results of classical rearrangement theory, where distances often correspond to simple graph structures (e.g. cycles (DCJ distance) or 2-cycles (breakpoint distance)) in specialized graphs, such as the breakpoint graph. The difference here is that the MBG can capture arbitrary (pangenome) graphs.

## 6 Evaluation

We implemented the SCJ-CARP reconstruction as well as utility functions to filter graph nodes based on size and extracting fixed size regions surrounding nodes from the graph. Relevant for the following experiments are the programs `carp`, which calculates the SCJ-CARP measure optionally filtering small nodes below a threshold and `carp-scan`, which calculates the SCJ-CARP measures for a fixed size region around each node. The implementation as well as snakemake [28] workflows for all following experiments are published on github^1^. The implementation is also available on bioconda under package name `carp`.

We evaluate our method for simulated datasets with regards to its ability to track the number of rearrangement events and to reconstruct ancestral adjacencies, and for its practical implications on biological datasets.

### 6.1 Simulated experiments

In order to evaluate how well SCJ-CARP quantifies rearragements, we used ZOMBI [13] to simulate phylogenies of genomes from an initial genome with 1000 markers. We used default parameters unless otherwise specified. By default, ZOMBI generates phylogenies with 20 extant genomes.

In order to simulate genomes of varying complexity, we scaled rates in ZOMBI for gene originations, duplications, losses, inversions and transpositions from 0.1 to 1.0 of the default rate. A comparison between this rate scale and the SCJ-CARP measure calculated is shown in Fig. 7 (A). We see that the SCJ-CARP measure tracks the number of simulated rearrangement operations well, particularly for low rates of rearrangements. In fact the Pearson and Spearman correlation coefficients for the entire experiment are 0.89 and 0.91 respectively, indicating a strong correlation between the number of simulated operations and the SCJ-CARP measure.

**Figure 7:**
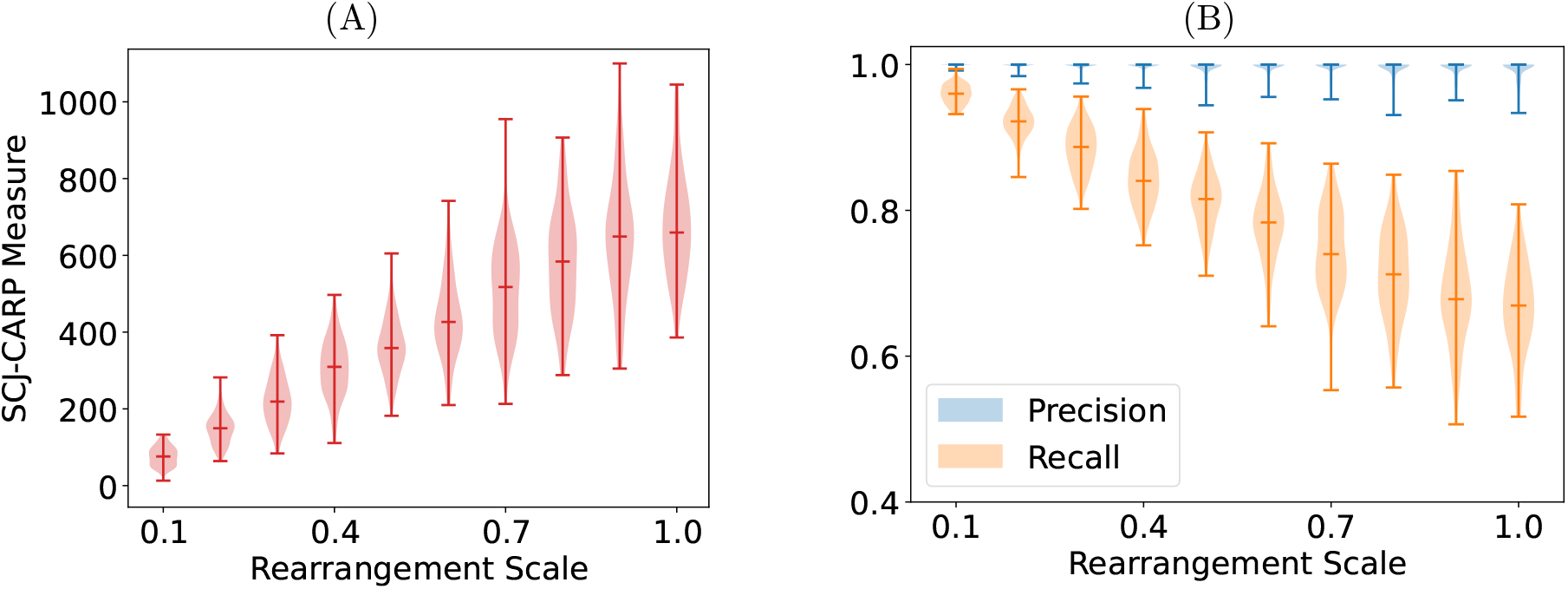
SCJ-CARP measure (A) and reconstruction precision and recall (B) with increasing numbers of rearrangements.

We then proceeded to evaluate in which way the marker adjacencies in the SCJ-CARP simplification correspond to the ancestral root simulated by ZOMBI. In this analysis, we limited ourselves to non-telomeric adjacencies, since in the SCJ model telomeres are best interpreted as uncertainty about which adjacency should be reconstructed, not as actual ends of chromosomes. We evaluated precision and recall using the adjacencies of the root genome as the ground truth. The results are shown in Fig. 7 (B). We see that the precision is high across all samples with the majority of samples in each step reaching 100% precision. However, recall degenerates quickly with increasing numbers of rearrangements. This is not suprising as the simplifications reconstructed by `carp` are extremely conservative – like in other problems under SCJ, ancestral genomes tend to fragment unless adjacencies are perfectly conserved.

### 6.2 Evaluating the SCJ-CARP measure of multiple pangenomes

We downloaded multiple genomes for 8 different taxa using NCBI datasets [36] to compare their structural complexity with SCJ-CARP. We aimed for 10 genomes with NCBI assembly level “complete” when possible and opted for “chromosome+” and a slightly reduced number of genomes when necessary. Details are found in Appendix A. We ran Minigraph-Cactus [23] and SibeliaZ [33] with two different settings to determine collinear blocks. We then ran `maf2synteny` [27] with default settings on the output of both tools.

We evaluated the SCJ-CARP measure on the resulting genome permutation files. Each run of `carp` took less than a second using four threads on a cluster machine. The calculated SCJ-CARP measures are visualized in Fig. 8.

**Figure 8:**
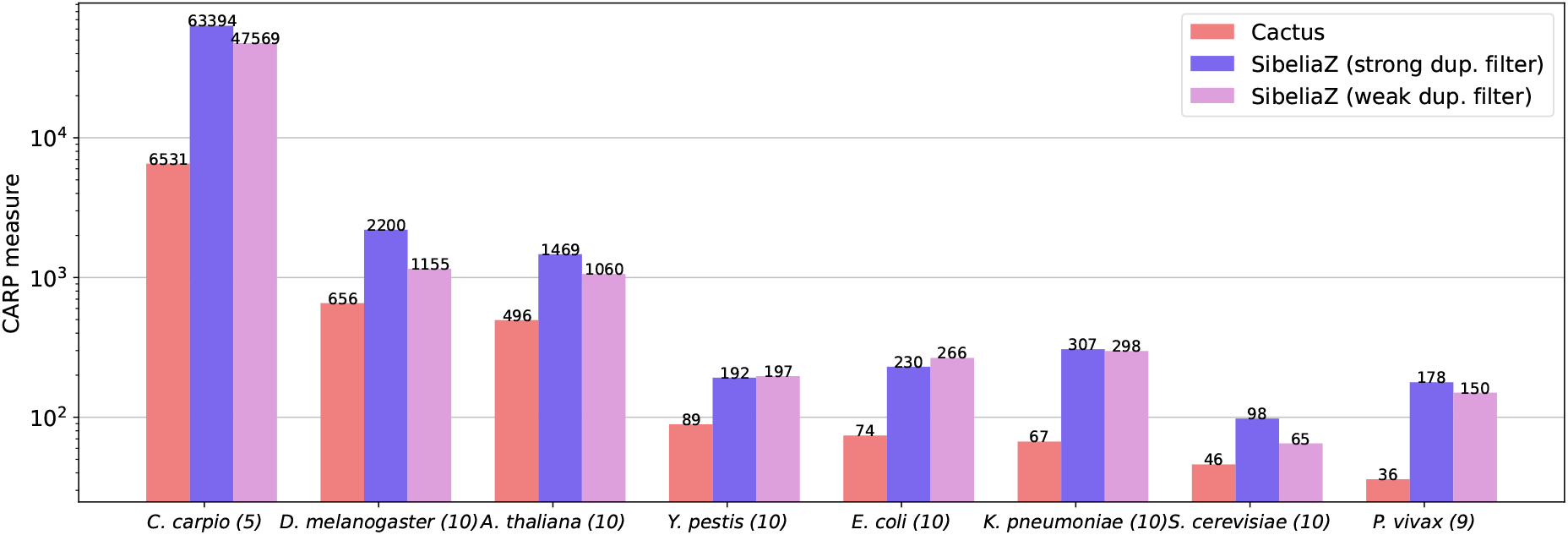
SCJ-CARP measures on pangenomes with collinear blocks detected by Minigraph-Cactus and SibeliaZ. Two different settings were used for SibeliaZ, a strong duplication filter (*a* = 2 × #genomes) and a weak duplication filter (*a* = 20 × #genomes). Shown in parentheses behind the taxon name is the number of genomes used as input to the block detection.

We see that the SCJ-CARP measures rank the relative pangenome complexity consistently even when using different block construction methods. The only disagreements are with respect to the order of *Y. pestis, E. coli* and *K. pneumoniae*, which are not well separated using any of the three block types, and the placement of *P. vivax*. Nonetheless, overall correlation between the CARP measures under different block definitions in this experiment is high (see Appendix Fig. 14).

Notably there are vast differences in the absolute SCJ-CARP measures between SibeliaZ and Minigraph-Cactus. Most interesting is the difference for *P. vivax*, for which both SibeliaZ runs evaluate to a much higher SCJ-CARP measure than Minigraph-Cactus, which yields the lowest SCJ-CARP measure overall. We conjecture that this is due to the differing definitions of blocks used by the two tools. It might also be the case that the SCJ model struggles to capture the rearrangements displayed by SibeliaZ. Another model may score the Minigraph-Cactus and SibeliaZ blocks more similarly. Thus, once solutions to CARP under more complex models are available, the differences between these segmentation approaches may become better explainable.

The discrepancy also emphasizes how crucial the initial block definition is for the comparison of genomes, a known issue in rearrangement analyses. Especially note that the absolute CARP measures on differently determined blocks are usually not comparable. Nonetheless, using blocks determined by the same method, SCJ-CARP gives a relatively coarse but quickly computable measure for the rearrangement complexity of a pangenome.

### 6.3 Comparing human pangenome graphs using CARP

Given the effect of different marker definitions on the CARP measures observed in the previous section, it stands to reason that the discrepancy between CARP measures of pangenome graphs constructed by different methods can give insights about how these graphs differ structurally. Notably, such structural differences have been observed between Minigraph, Minigraph-Cactus and PGGB during the construction of the first human pangenome reference [31]. There the authors note that Minigraph-Cactus includes smaller variants than Minigraph while still constructing similar nodes for duplicate regions, while PGGB’s all-vs-all alignment strategy creates a graph that collapses duplications resulting in an overall more complex graph (see Extended Data Fig. 3 in [31] for an example).

We downloaded the three respective graphs from the human-pangenomics S3 bucket using the CHM13-version of the Minigraph and Minigraph-Cactus graphs. We proceeded to analyze the graphs using `carp`, filtering out nodes below a threshold of progressively larger size. We measured the number of nodes in the graph and the SCJ-CARP measure. We show these results as well as the SCJ-CARP measure relative to the number of nodes in Fig. 9.

**Figure 9:**
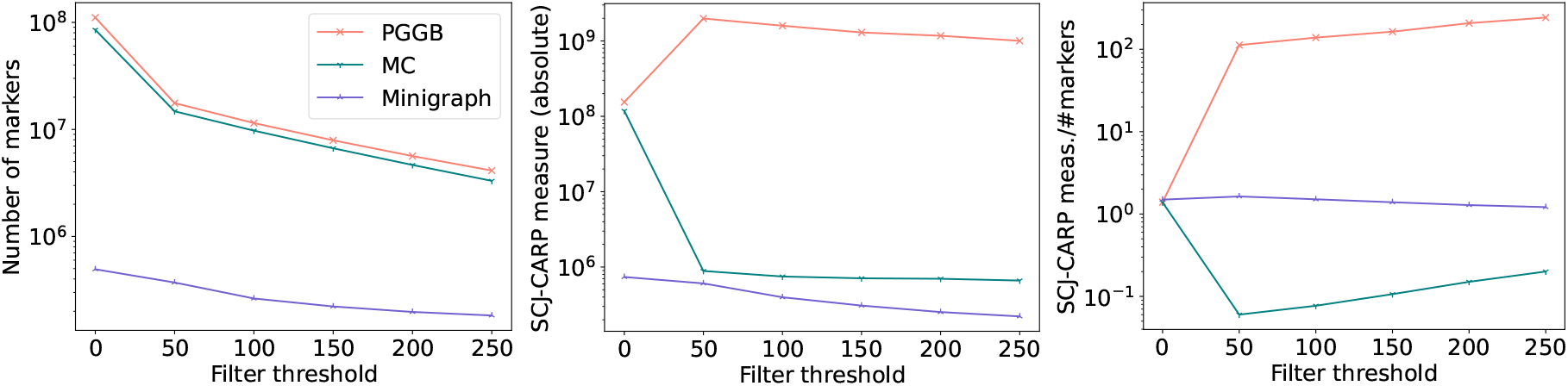
CARP statistics for various levels of node filtering on human pangenome graphs built by PGGB, Minigraph-Cactus (MC) and Minigraph from [31]. Nodes of a size smaller than the filter threshold were removed and each path bridging two nodes that pass the threshold consisting of only filtered nodes was replaced with a single edge.

Without filtering the nodes (Filter threshold 0), we see that Minigraph is an outlier, while Minigraph-Cactus and PGGB are close together, both in the number of nodes and in the absolute SCJ-CARP measure. Nonetheless, the relative SCJ-CARP measure of all three tools is similar. Progressively filtering nodes, differences between Minigraph, Minigraph-Cactus and PGGB become more apparent. While the number of nodes decreases for all three graphs, again with a similar trajectory for Minigraph-Cactus and PGGB, the SCJ-CARP measure shows that PGGB’s complexity increases, while the complexity of Minigraph-Cactus and Minigraph is now much more similar.

From these results one can conclude that while the graphs of PGGB and Minigraph-Cactus have similar numbers of nodes, the way variants are represented in Minigraph and Minigraph-Cactus is much more similar to each other than to PGGB. PGGB in turn tends to produce highly connected small nodes that lead to an increase in connectedness and consequently higher complexity once they are filtered.

These observations are of course consistent with the knowledge that Minigraph-Cactus graphs are constructed by adding smaller variants to Minigraph graphs while PGGB follows a different, alignment-based paradigm. However, note that this knowledge of the graph construction methods is not necessary to interpret the results computed by SCJ-CARP.

On 16 threads, calculations for Minigraph were completed within seconds while calculations for Minigraph-Cactus finished in a matter of minutes. The longest calculation for PGGB completed after about four hours. Overall, the entire analysis was completed within 20 hours. We give all runtimes of SCJ-CARP as Appendix Fig. 15.

We conclude that SCJ-CARP allows an analysis of pangenome graphs that highlights their differing structural qualities without prior knowledge of how these graphs were constructed. Additionally, the time frame for such an analysis is reasonable, taking about one day for human sized pangenomes.

## 7 Discussion

We developed CARP, a new rearrangement problem designed to quantify the rearrangement complexity of pangenomes. This problem relates intuitively to both pangenome graphs and classical genome rearrangement, thus forming a first step towards a joint modeling of these two, highly connected areas. We presented a solution to CARP under the simple SCJ model. This solution already enables fast analyses of pangenomes and their graphs, which relate valuable insights about construction methods and structural complexity to a formal rearrangement perspective. We expect this potential to further unfold once solutions under other models become available. For example, one could characterize a pangenome by a vector of CARP measures of different models (i.e. inversions, block interchanges, etc.), indicating which model best explains its complexity. This would also allow a high level comparison between pangenomes that do not share any sequence material. A prerequisite to perform such a comparison is some normalization of the CARP measure. Like other rearrangement-based methods, CARP yields an absolute number of rearrangements, which of course depends on the size and number of the input genomes. Future research may explore normalizing the CARP measure as has been done with other pangenome metrics (see for example [37]).

We have also seen that the CARP measure under the SCJ-model corresponds to a simple to evaluate graph metric in a general pangenome graph, which mirrors the fact that this is the case for many models in classical rearrangement analyses in the specialized breakpoint graph. It thus stands to reason that the CARP measure for other rearrangement models may be related to other known graph theoretic quantities. This would allow the study of abstract graph properties under the more biologically meaningful lens of rearrangement models and, vice versa, the mathematical understanding of a rearrangement model would be enhanced by connecting it to graph theory in general.

Furthermore, it is worth investigating the implications that Path Altering Operations used in a given scenario make about the phylogenetic relationship between genomes of the pangenome. This may well lead to an easily computable rearrangement-based tree reconstruction.

## Acknowledgements

This work was funded by the Federal Ministry of Research, Technology and Space in the frame of de.NBI & ELIXIR-DE (W-de.NBI-004).

LB has received funding from the European Union’s Horizon 2020 research and innovation programme under the Marie Skłodowska-Curie grant agreement PANGAIA No. 872539.

This work was supported by the de.NBI Cloud within the German Network for Bioinformatics Infrastructure (de.NBI) and ELIXIR-DE (Forschungszentrum Jülich and W-de.NBI-001, W-de.NBI-004, W-de.NBI-008, W-de.NBI-010, W-de.NBI-013, W-de.NBI-014, W-de.NBI-016, W-de.NBI-022).

## Appendix

### A Pangenome comparison – datasets

### B Scoring region complexities in a *Bacillus subtilis* de Bruijn graph

Typically the fastest way to build a pangenome graph is to construct a de Bruijn graph of all genome sequences in the pangenome of interest. However, de Bruijn graphs tend to be large and hard to interpret, especially in regions with a high complexity (“hair balls”), which typically arise due to nodes that appear very frequently in the pangenome. The SCJ-CARP measure can be used to score the complexity of the subgraph surrounding a node.

To demonstrate this possibility, we downloaded all 1508 available *Bacillus subtilis* genomes in genbank using NCBI datasets [36] and constructed a de Bruijn graph using Bifrost [26] for *k* = 31. The resulting graph had a total number of 3, 665, 906 nodes. We then used the `carp-scan` program to extract regions of 300 BP around a node and to score this environment using the SCJ-CARP measure. On 8 threads on a cluster machine, `carp-scan` finished after 8 minutes and 4 seconds. We show the histogram of SCJ-CARP measures of the 3, 629, 054 least complex nodes and examples of node environments in Fig. 10.

**Figure 10:**
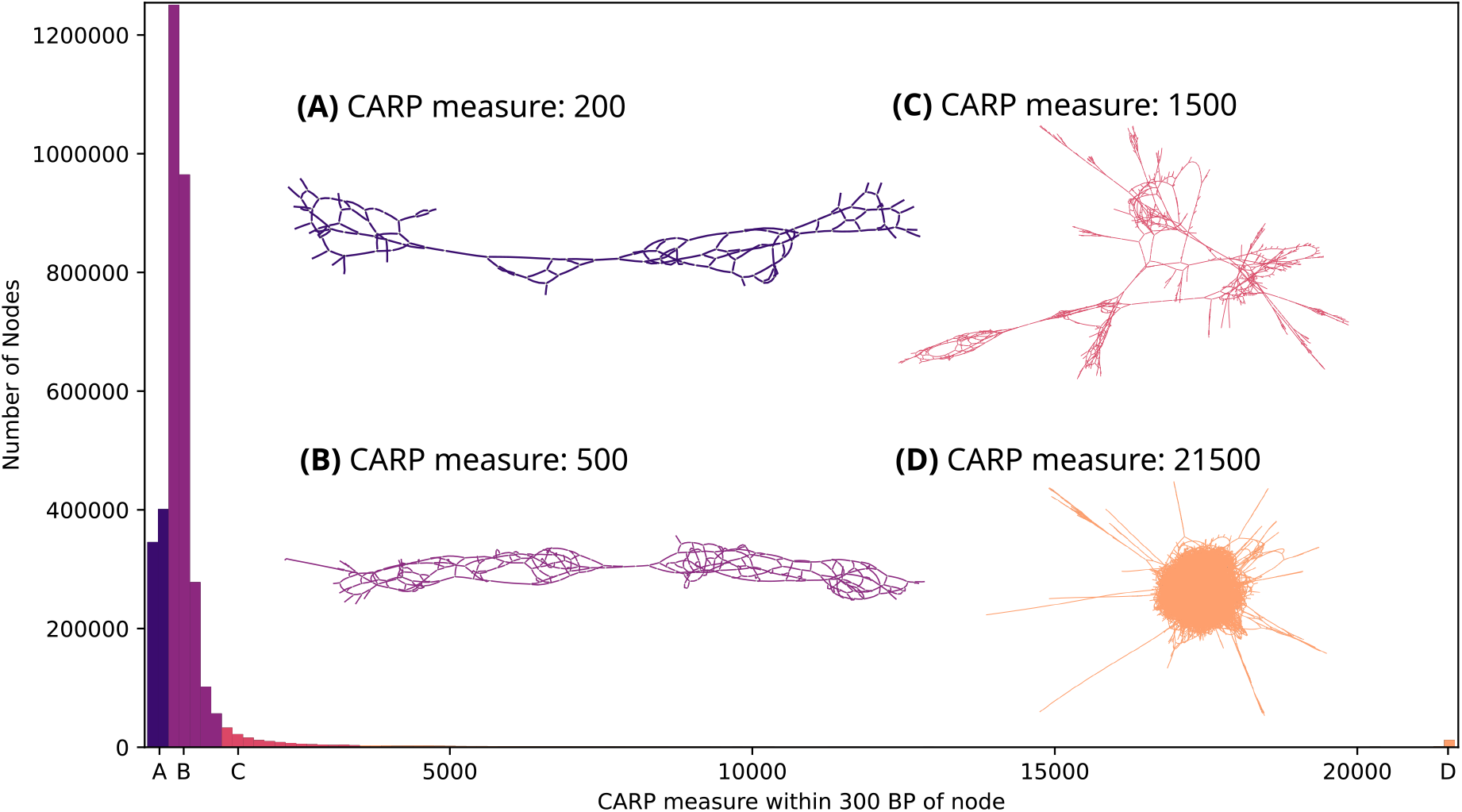
Histogram of SCJ-CARP measures in the 300 BP environment of nodes in a compacted de Bruijn graph built from 1508 *Bacillus subtilis* genomes. For a better visualization, the histogram is artificially cut after 21500; the highest SCJ-CARP measure of a node environment was 175702. (A) to (D) visualize example environments from distinct regions of the distribution. These visualizations were created with Bandage [48].

We see that the majority of nodes are surrounded by a subgraph with a CARP measure below 1500, which appear to be interpretable (see Examples (A), (B) and (C) in Fig. 10), although ease of interpretation decreases as the complexity identified by the SCJ-CARP measure increases. True “hair ball” subgraphs, such as shown as Example (D) in Fig. 10, seem to be the exception in this graph, but there is a cluster of these nodes around SCJ-CARP measure 21500.

We also performed additional analyses varying the environment size. These histograms can be found in Appendix Fig. 12. Moreover, the `carp-scan` program can color graph nodes based on their surrounding SCJ-CARP measures, which enables to easily visualize where in the graph complexity arises. We give an example in Appendix Fig. 13. Additionally, one can map these complexities back to a reference genome and visualize it with a genome browser. We give an example in Fig. 11 using the UCSC Genome Browser [39].

**Figure 11:**
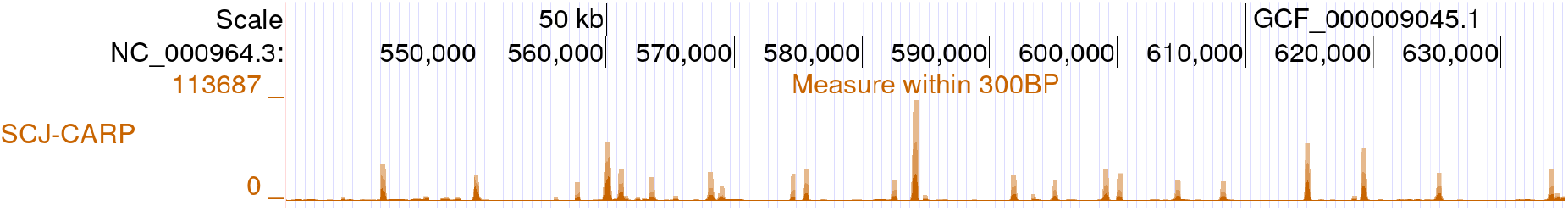
300 BP environment complexities in the de Bruijn graph (*k* = 31) mapped to the *Bacillus subtilis* reference `GCF_000009045.1`. Shown is the region `NC_000964.3:534,995-635,004` in the UCSC genome browser [39].

**Figure 12:**
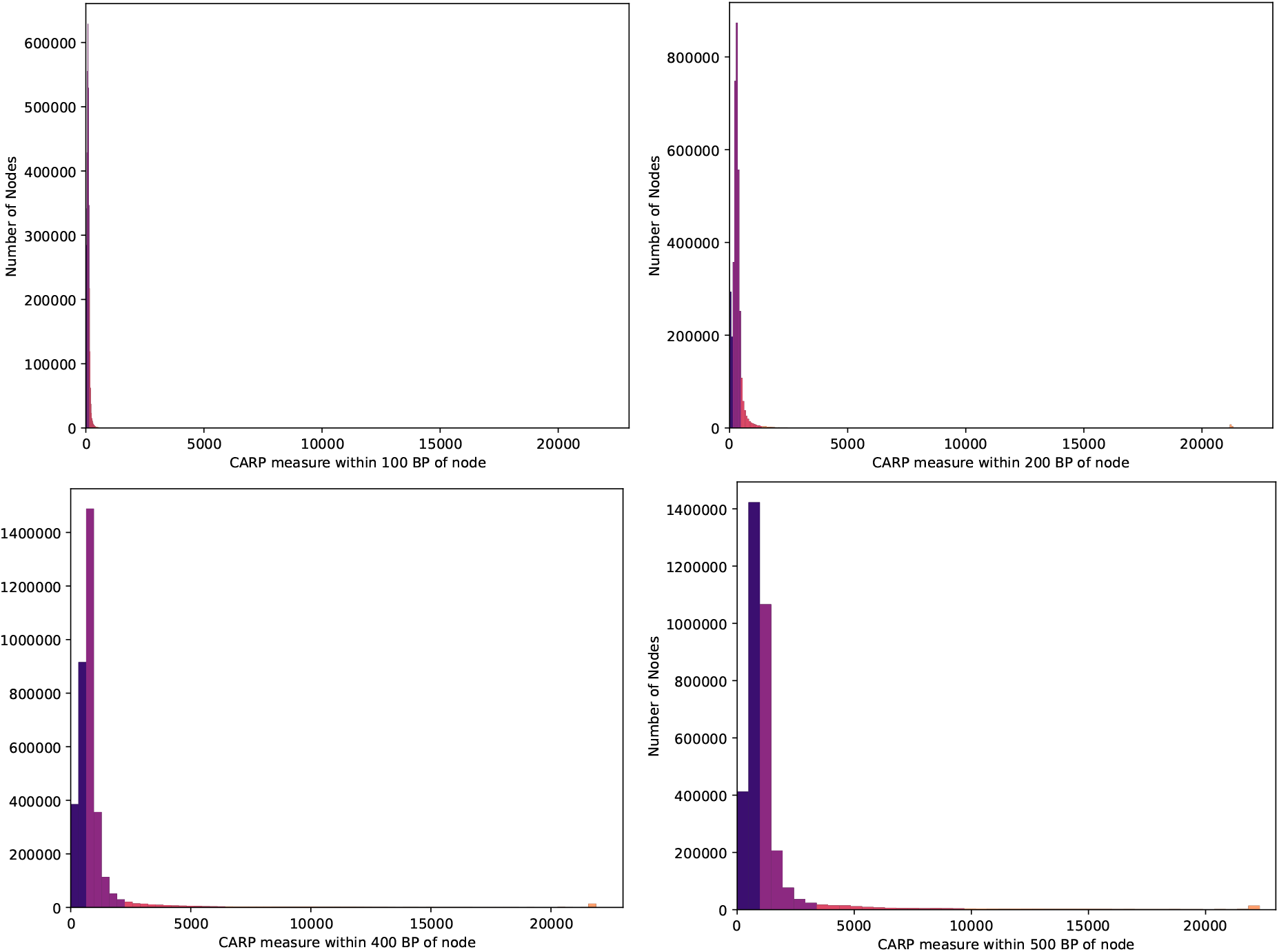
Alternative histograms for the *Bacillus subtilis* dataset with node environments 100, 200, 400 and 500. Shown is the same excerpt of the histogram as in Fig. 10.

**Figure 13:**
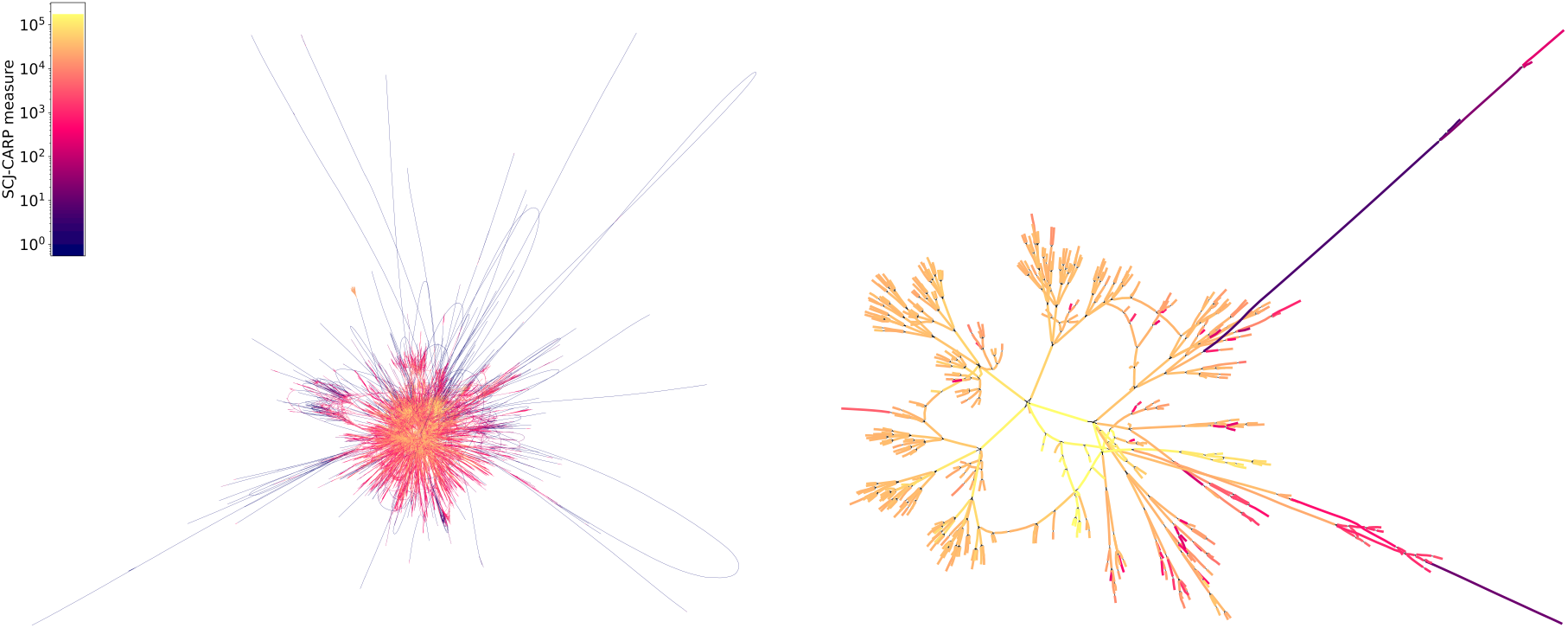
Coloring of nodes in the de Bruijn graph (*k* = 31) of 1508 *Bacillus subtilis* genomes by SCJ-CARP measure of surrounding 300 BP. Shown is a 20-node (left) and 6-node environment around the most complex node visualized with Bandage [48]. Note that the color scheme is not the same as in the histograms (Figures 10 and 12).

**Figure 14:**
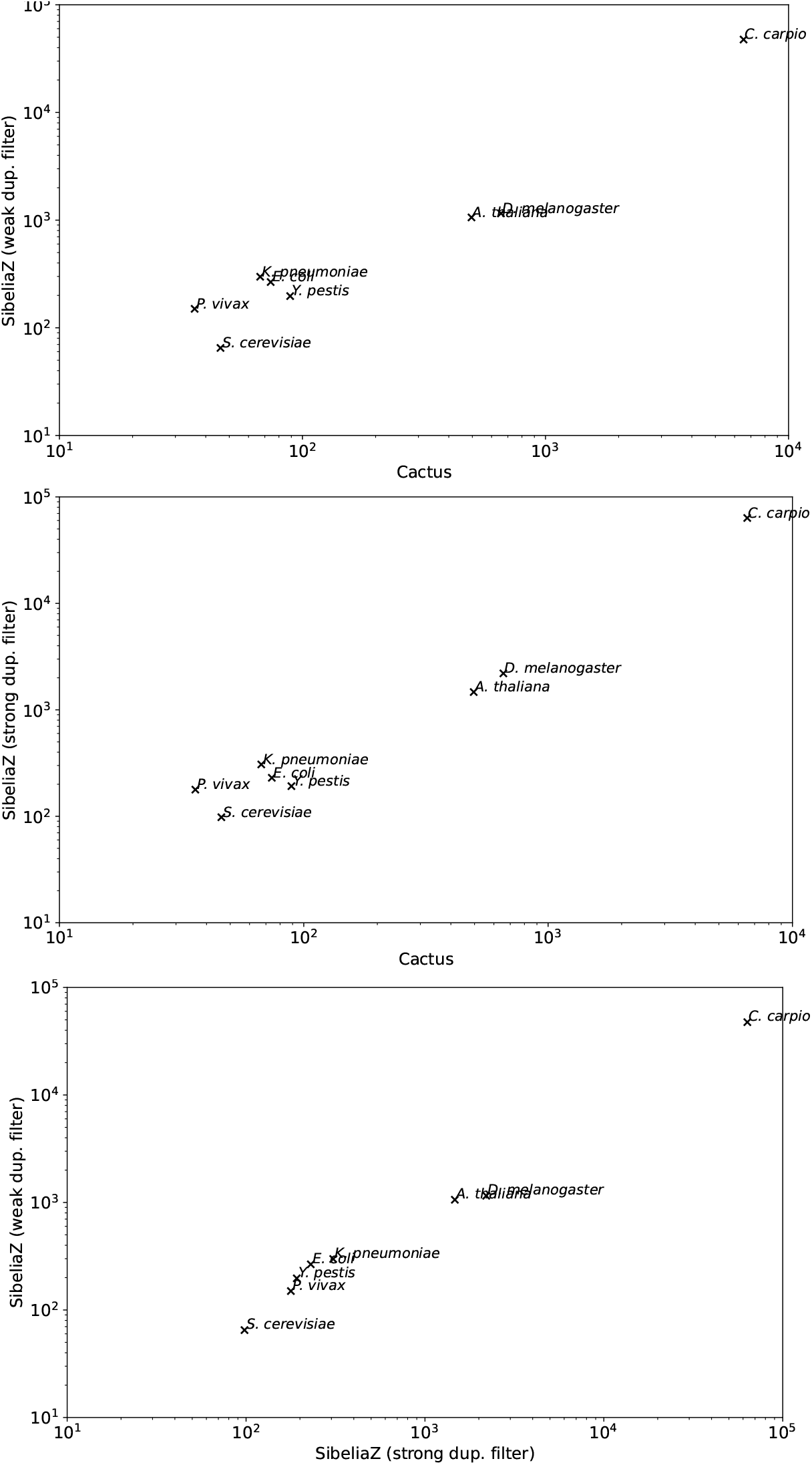
Pairwise scatterplots of SCJ-CARP measures calculated on different block types as a log-log plot.

**Figure 15:**
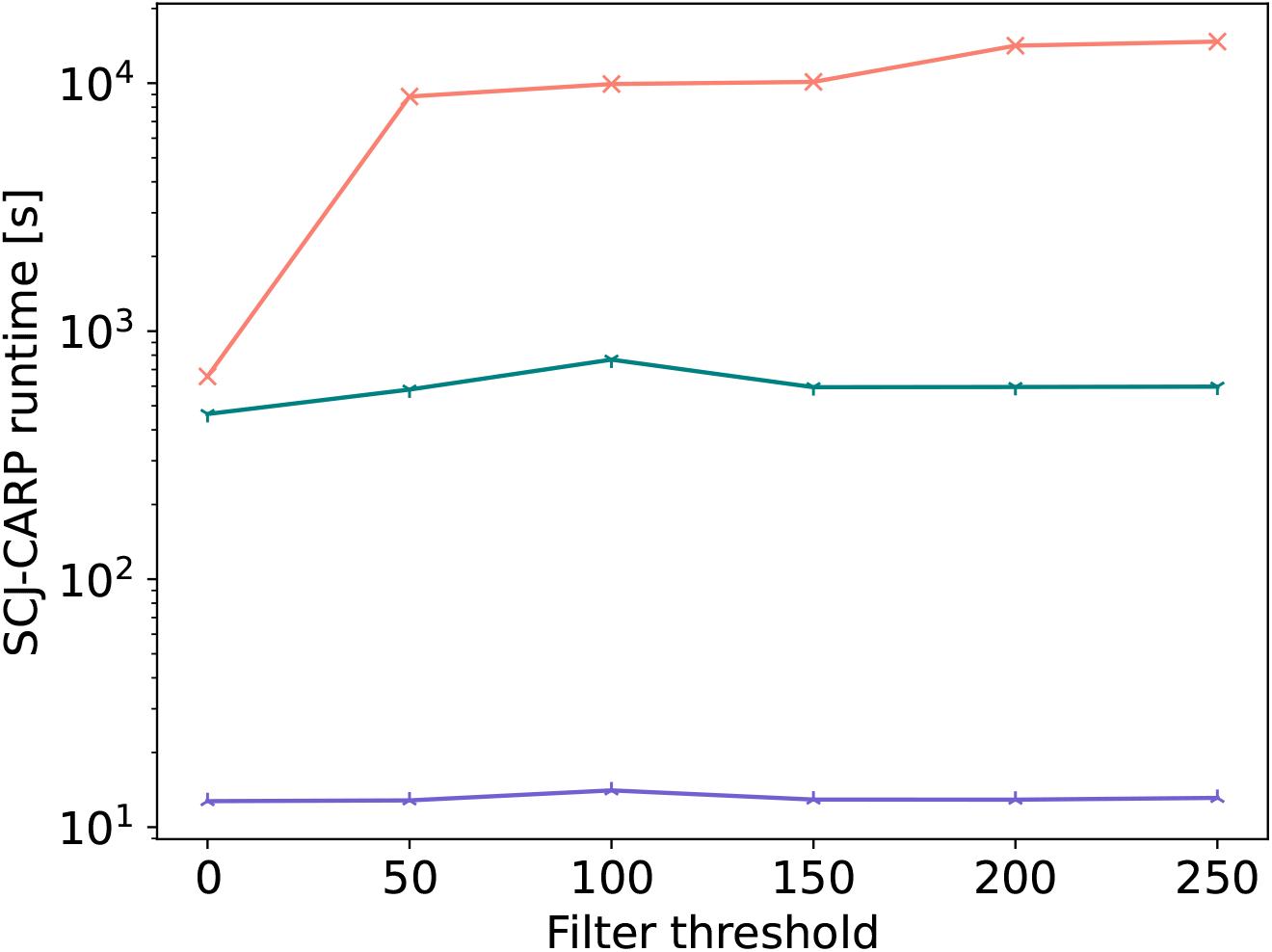
CARP runtime for various levels of node filtering on human pangenome graphs built by PGGB, Minigraph-Cactus (MC) and Minigraph from [31]. Nodes of a size smaller than the filter threshold were removed and each path bridging two nodes that pass the threshold consisting only of filtered nodes was replaced with a single edge. The SCJ-CARP software was run with 16 threads on a cluster machine.

One can thus use SCJ-CARP to identify regions in a pangenome graph that are already interpretable and regions that still need refinement, which could be a useful tool or evaluation metric for pangenome graph construction methods. Moreover, this allows in principle to compare pangenomes and examine at which level structural complexity arises and how it is distributed throughout the pangenome graph.

### C Supplementary Figures

## Footnotes

1 https://github.com/gi-bielefeld/scj-carp

